# MPAC: a computational framework for inferring pathway activities from multi-omic data

**DOI:** 10.1101/2024.06.15.599113

**Authors:** Peng Liu, David Page, Paul Ahlquist, Irene M. Ong, Anthony Gitter

## Abstract

Fully capturing cellular state requires examining genomic, epigenomic, transcriptomic, proteomic, and other assays for a biological sample and comprehensive computational modeling to reason with the complex and sometimes conflicting measurements. Modeling these so-called multi-omic data is especially beneficial in disease analysis, where observations across omic data types may reveal unexpected patient groupings and inform clinical outcomes and treatments. We present Multi-omic Pathway Analysis of Cells (MPAC), a computational framework that interprets multi-omic data through prior knowledge from biological pathways. MPAC leverages network relationships encoded in pathways through a factor graph to infer consensus activity levels for proteins and associated pathway entities from multi-omic data, runs permutation testing to eliminate spurious activity predictions, and groups biological samples by pathway activities to allow identifying and prioritizing proteins with potential clinical relevance, e.g., associated with patient prognosis. Using DNA copy number alteration and RNA-seq data from head and neck squamous cell carcinoma patients from The Cancer Genome Atlas as an example, we demonstrate that MPAC predicts a patient subgroup related to immune responses not identified by analysis with either input omic data type alone. Key proteins identified via this subgroup have pathway activities related to clinical outcome as well as immune cell compositions. Our MPAC R package, available at https://bioconductor.org/packages/MPAC, enables similar multi-omic analyses on new datasets.

## Introduction

Cancer is a complex set of diseases with a great diversity of genomic aberrations and altered signaling pathways (Hanahan, 2022). The Cancer Genome Atlas (TCGA) generated data spanning copy number alteration (CNA), DNA mutation, DNA methylation, mRNA expression, microRNA expression, and protein expression for thousands of tumor samples, leading to many insights into the cancers that were profiled (Hoadley *et al*., 2018). In addition, this extensive multi-omic data provides clues to tumor regulation, which have led to the development of many computational methods to integrate multi-omic data to obtain comprehensive views on cancer (Picard *et al*., 2021; Maghsoudi *et al*., 2022; G. L. Stein-O’Brien *et al*., 2018).

In particular, biological pathway-based approaches have been demonstrated as a powerful way to integrate multi-omic data (reviewed in Maghsoudi et al. 2022). Altered expression or function of different genes in the same pathway can have similar impacts on overall pathway activity. Similarly, diverse alterations of expression or function of the same gene or its protein product—e.g. through DNA mutations, CNAs, or changes in epigenetic modifications, transcript expression, or protein translation, stability, or post-translational modifications—can also suppress, stimulate, or otherwise modulate a particular pathway. These properties allow modeling based on multi-omic inputs to infer pathway activity to more accurately reflect underlying biology than modeling based on a narrow, incomplete view from a single genomic data type. Accordingly, whereas a single data type rarely contains the full explanation for oncogenesis, pathway-based approaches are a particularly advantageous way to understand cancer mechanisms.

Several notable pathway-based methods have demonstrated the benefits of multi-omic data integration for cancer interpretation. For example, Multi-omics Master-Regulator Analysis (MOMA) identified 112 distinct tumor subtypes and 24 conserved master regulator blocks across 20 TCGA cohorts (Paull *et al*., 2021). OncoSig delineated tumor-specific molecular interaction signaling maps for the full repertoire of 715 proteins in the COSMIC Cancer Gene Census (Broyde *et al*., 2021). COSMOS combined signaling, metabolic, and gene regulatory networks to capture crosstalks within and between multi-omics data (Dugourd *et al*., 2021). PAthway Recognition Algorithm using Data Integration on Genomic Models (PARADIGM) integrates multi-omic data via a factor graph to infer activities of all the proteins in a pathway network (Sedgewick *et al*., 2013; Vaske *et al*., 2010). Initially, PARADIGM was successfully applied to breast cancer and glioblastoma patients using CNA and gene expression microarray data to find clinically relevant groups and associated pathways. It was further applied to reveal multiple low-frequency, but high-impact mutations in glioblastoma, ovarian, and lung cancers (Ng *et al*., 2012) and was incorporated into the standard TCGA analysis pipeline (Hoadley *et al*., 2018, 2014).

Despite such successes, there are still opportunities to further improve multi-omic modeling. MOMA and OncoSig focused on direct interactions around master regulators for transcription. The indirect effects of proteins further downstream of the master regulators in biological pathways were not considered. PARADIGM’s application across many cancer types focused on grouping patients by their inferred pathway levels or enriched pathways (Hoadley *et al*., 2018; Berger *et al*., 2018). But in-depth analysis on the molecular basis of patient grouping, careful interpretation of its inferred pathway levels, and an end-to-end computational process are lacking. PARADIGM’s inferred pathway levels are abstract quantities indicating the log-likelihood ratio of proteins being activated or repressed, but they represent neither protein abundance nor any particular post-translational modification and cannot be experimentally measured. Other existing patient stratification methods by multi-omic data either do not use biological pathway information (Duan *et al*., 2021) or rely on unrealistically small pathways (Zhao *et al*., 2021). As a result, it is hard to identify key proteins from a broad perspective with meaningful biological interpretation and clinical implication.

Here, we develop a computational framework, named Multi-omic Pathway Analysis of Cells (MPAC), to integrate multi-omic data for understanding cellular networks. It is built upon the PARADIGM method with notable improvements including providing enhanced insights to the molecular basis and clinical implications of pathway-based patient groups as well as streamlining the whole computational process. In this work, we apply MPAC to Head and Neck Squamous Cell Carcinoma (HNSCC), which accounts for ∼500,000 deaths per year worldwide (Mody *et al*., 2021). First, we describe how MPAC improves upon PARADIGM. Next, we apply MPAC to TCGA HNSCC data and group patients by their significantly altered pathways. Among other results, MPAC predicts a patient group that is enriched with immune response pathways, and this group cannot be predicted from the individual omic data types alone. Investigating this group identifies seven proteins that have activated pathway levels associated with better overall survival. These findings are validated by a holdout set of TCGA HNSCC samples. We demonstrate MPAC’s improvements over PARADIGM by showing that PARADIGM cannot identify such an immune response group. We also evaluate MPAC’s robustness by running it with different settings and another TCGA cancer type, cholangiocarcinoma. Lastly, we present an interactive R Shiny app that lets users explore all the results generated from this work.

## Results

### An overview of MPAC and its improvements upon PARADIGM

We developed MPAC to integrate multi-omic data to identify key pathways and proteins with biological and clinical implications, and to predict new patient groups associated with distinct pathway alterations. MPAC’s workflow contains eight steps (Figure 1 and Supplementary Note 1; see also Methods for more details): (Step 1) From CNA and RNA-seq data, determine genes’ CNA and RNA ternary states (i.e., repressed, normal, or activated). CNA and RNA-seq data are selected as the input multi-omic data because PARADIGM had shown success with them (Vaske et al. 2010; Sedgewick et al. 2013; Hoadley et al. 2018); (Step 2) Use CNA and RNA state together with a comprehensive biological pathway network from TCGA (Hoadley *et al*., 2018) to calculate pathway levels with PARADIGM’s factor graph model. The TCGA pathway network characterizes interactions at both transcriptional and post-transcriptional levels. PARADIGM’s inferred pathway levels are calculated not only for proteins, but also for several other types of pathway entities, such as protein complexes and gene families; (Step 3) Permute input CNA and RNA data for filtering inferred pathway level in the next step. CNA and RNA states are permuted randomly between genes in each patient. Inferred pathway levels for each pathway entity are calculated with PARADIGM from 100 sets of permuted data to build a background distribution representing inferred pathway levels observed by chance; (Step 4) Inferred pathway levels computed from real data are compared with those from permuted data to filter out inferred pathway level changes that could be observed by chance. Because both the real and permuted pathway levels are different between patients, this filtering step creates a patient-specific set of inferred pathway levels representing each patient’s unique pathway alteration profiles; (Step 5) From the remaining pathway networks, retain the connected component that contains the largest number of connected pathway entities. This focuses on the main subset of entities that are connected in the pathway network and presumably lead to similar functional alterations; (Step 6) Build patient pathway profiles and predict patient groups. Each patient’s pathway profile contains a selected set of 2,085 cancer- and general biology-relevant Gene Ontology (GO) terms (Gene Ontology Consortium *et al*., 2023; Kuenzi *et al*., 2020). Each GO term is characterized by an over-representation test p-value on entities selected from the previous step. P-values for all the GO terms are adjusted for multiple hypothesis testing and log-transformed to group patients (Lun *et al*., 2016); (Step 7) Identify key proteins that all have activated or repressed inferred pathway levels between patients from the same group; (Step 8) Compare data on key proteins with patients’ clinical data to evaluate potential impact of protein inferred pathway levels on patients’ clinical outcomes.

**Figure 1.**
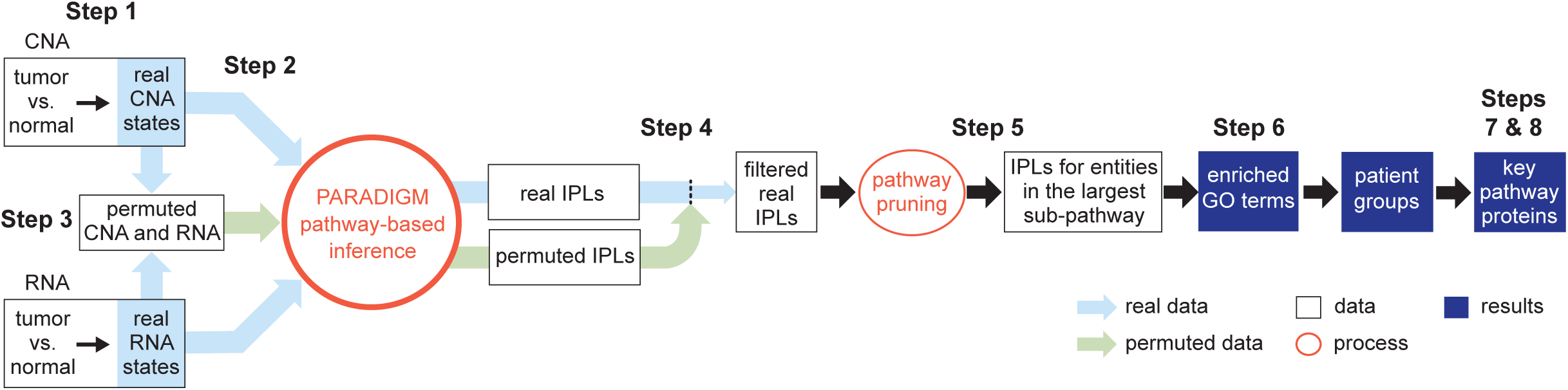
Overview of the MPAC workflow. MPAC calculates inferred pathway levels (IPLs) from real and permuted CNA and RNA data. It filters real IPLs using the permuted IPLs to remove spurious IPLs. Then, MPAC focuses on the largest pathway subset network with filtered IPLs to compute Gene Ontology (GO) term enrichment, predict patient groups, and identify key group-specific proteins.

MPAC makes multiple improvements (Supplementary Table 1) over PARADIGM (Vaske *et al*., 2010), which MPAC runs as a subroutine in Steps 2 and 3. PARADIGM simply divided all the genes into three states with an equal number of entries. In contrast, in Step 1, MPAC defines a gene’s RNA state as normal, activated or repressed for each patient by testing the level of the relevant RNA in that tumor sample for a significant increase or decrease (two standard deviations from the mean of a Gaussian distribution, which equals the commonly used *p* < 0.05 cutoff) of that RNA’s expression distribution in normal tissue samples. In Steps 3 and 4, as noted above, MPAC filters pathway entities for significant inferred pathway level differences from randomly permuted input. Although some PARADIGM applications also used permutations, permutations were not implemented as part of the software, nor were their results used for downstream analysis (Vaske *et al*., 2010; Sedgewick *et al*., 2013). In Step 5, MPAC focuses on the largest patient-specific pathway network subset. This improvement removed from consideration entities in tiny pathways, which were assumed to have less impact on patient pathway alterations and may contribute more noise than signal when predicting patient groups. In Steps 6–8, MPAC provides downstream analysis functions to define patient pathway alterations, predict patient groups, and identify key proteins with potential clinical implications. MPAC is available as an R package on Bioconductor (https://bioconductor.org/packages/MPAC) to streamline the whole process from preparing the omic input data to identifying key proteins for a patient group.

### MPAC predicted an immune response HNSCC group not found by CNA or RNA-seq data alone

We applied MPAC to TCGA HNSCC patients to predict patient groups by their pathway alterations. We selected the 492 patients that had CNA, RNA-seq, and overall survival data available. Of these 492 tumors, 89 carried human papillomavirus DNA (HPV+) and 403 did not (HPV-), a distinction linked to major differences in HNSCC tumor biology and clinical treatment response (Powell *et al*., 2021). HPV+ HNSCC is mainly caused by HPV’s E6 and E7 proteins, whereas HPV- HNSCC has much higher mutation loads and distinct oncogenic pathways than HPV+. Because of such differences, we applied MPAC to the two HPV subtypes separately. We further randomly divided patients into exploratory sets (71 HPV+ and 322 HPV-) and validation sets (18 HPV+ and 81 HPV-) (Supplementary Table 2). Our goal was to first tune MPAC and identify pathway-based patient groups in the exploratory set and then test our discoveries in the validation set. MPAC identified five groups from each HPV subtype based on the patient pathway profiles (Figure 2 and Supplementary Figure 1A). For HPV+ patients (Figure 2), four of the five groups had distinct pathway features. Group I patients had alterations mainly in immune response pathways, groups II and IV in cell cycle pathways, and group V in morphogenesis pathways. Group III had pathway alterations in some patients but did not show an obvious biologically meaningful consensus profile. For HPV- patients (Supplementary Figure 1A), three of the five patient groups had distinct pathway alterations: groups I and IV in cell cycle pathways and group III in immune response pathways. Groups II and V did not show obvious consensus pathway features. The distinct pathway features for many of the patient groups suggested that MPAC is capable of building biomedically relevant patient pathway profiles and predicting patient groups.

**Figure 2.**
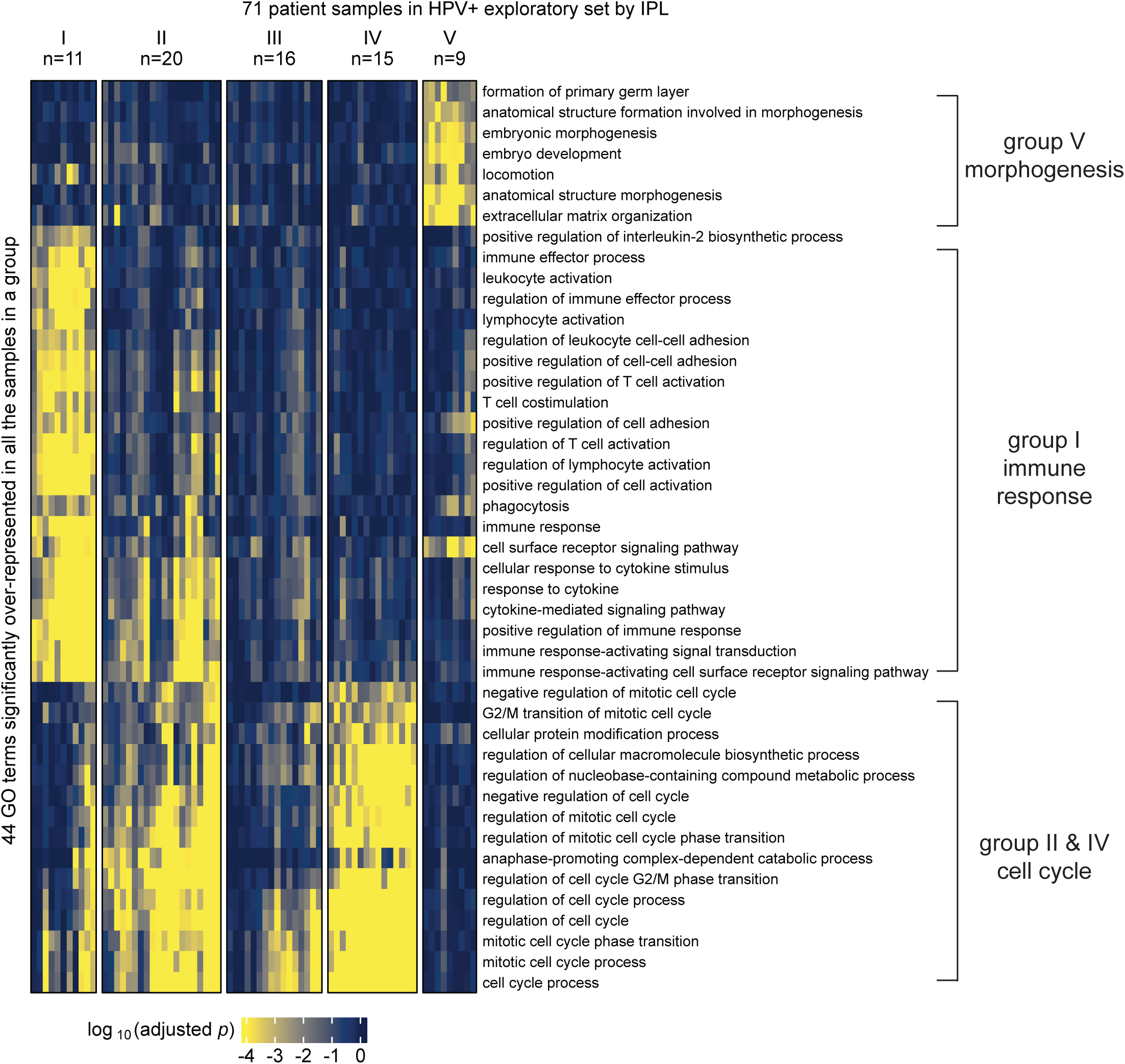
MPAC predicted functionally distinct patient groups in the HPV+ exploratory set. Patient groups were derived from GO term enrichment based on inferred pathway levels (IPLs).

MPAC predicted the above patient groups with distinct pathway profiles by integrating multi-omic data. We found that the same groups and pathways cannot be found by examining individual omic data types alone. Starting from either CNA or RNA-seq data, we conducted two tests: one performing GO enrichment on each single omic data type and then grouping enriched GO terms, like the MPAC workflow, and the other by grouping patients via their single omic data first and then finding commonly enriched GO terms within each group. In the first test, HPV+ patients’ CNA data had very few GO terms enriched, and even these were only enriched in a small number of patients (Supplementary Figure 2A). As a result, no group prediction could be made. RNA-seq data was more informative than CNA data, and four patient groups could be predicted (Supplementary Figure 2B). Groups III and IV were related to cell cycle and morphogenesis pathways, respectively, both of which had also been observed in the MPAC results. The immune response patient group predicted by MPAC, however, was not observed from CNA or RNA-seq data, indicating a unique insight from MPAC. For HPV- patients, CNA data did not lead to any patient groups due to insufficient significantly enriched GO terms (Supplementary Figure 1B). RNA-seq data led to six groups. Two of them were related to cell cycle and immune response (Supplementary Figure 1C), which were also observed in MPAC results.

To demonstrate the robustness of this result, we performed another test by grouping patients first and followed by GO enrichment. We applied K-means clustering to the RNA-seq data and divided HPV+ patients into two to six groups. The cluster membership remained stable under different numbers of groups (Supplementary Figure 3A). Therefore, we used five groups (Supplementary Figure 3B) for GO enrichment analysis so that every group had at least two samples while maintaining as many groups as possible. Groups I, II, IV, and V did not have any top GO terms related to immune response (Supplementary Figure 3 C–D and F–G). Group III was predominantly enriched with cell cycle-related GO terms with only one GO term (lymphocyte activation) related to immune response (Supplementary Figure 3E). Moreover, this single term was less consistently enriched than the >20 immune response GO terms from MPAC (Figure 2A). We performed the same analysis on CNA data. Stable grouping membership was observed (Supplementary Figure 4A), and the three-groups result (Supplementary Figure 4B) was used for GO enrichment analysis. No GO term was significantly overrepresented in at least half of the samples in any group. In summary, by jointly modeling both CNA and RNA-seq data, MPAC identified a large and unique HPV+ patient group related to immune response that could not be recovered from either individual omic data type alone.

### Proteins from the HPV+ immune response group associated with patient overall survival

Given that MPAC discovered an immune response patient group that could not be found by CNA or RNA-seq data alone, we were interested in pathway submodules and key proteins shared by the eleven patients in this group. We defined a pathway submodule as a pathway subset containing ≥ 5 entities, at least one of which was a protein with input omic data. We required that all submodule entities must have activated or repressed inferred pathway levels in the eleven patients. Four such submodules were identified (Figure 3A). They contained five to twelve entities and collectively eight proteins (Figure 3A, red ovals). Seven of these proteins, CD28, CD86, TYK2, IL12RB1, LCP2, FASLG, and CD247, had activated inferred pathway levels in all eleven group I patients (Figure 3B), suggesting a consensus functional role across patients within this immune response group. Interestingly, prior studies collectively showed that gene expression levels of the seven proteins except for LCP2 are associated with immune infiltration in HNSCC (Zhu *et al*., 2022; Wang *et al*., 2022; He *et al*., 2022; Chi *et al*., 2022; Chen *et al*., 2022; de Vos *et al*., 2020). Below we show a similar association for the patient group analyzed here.

**Figure 3.**
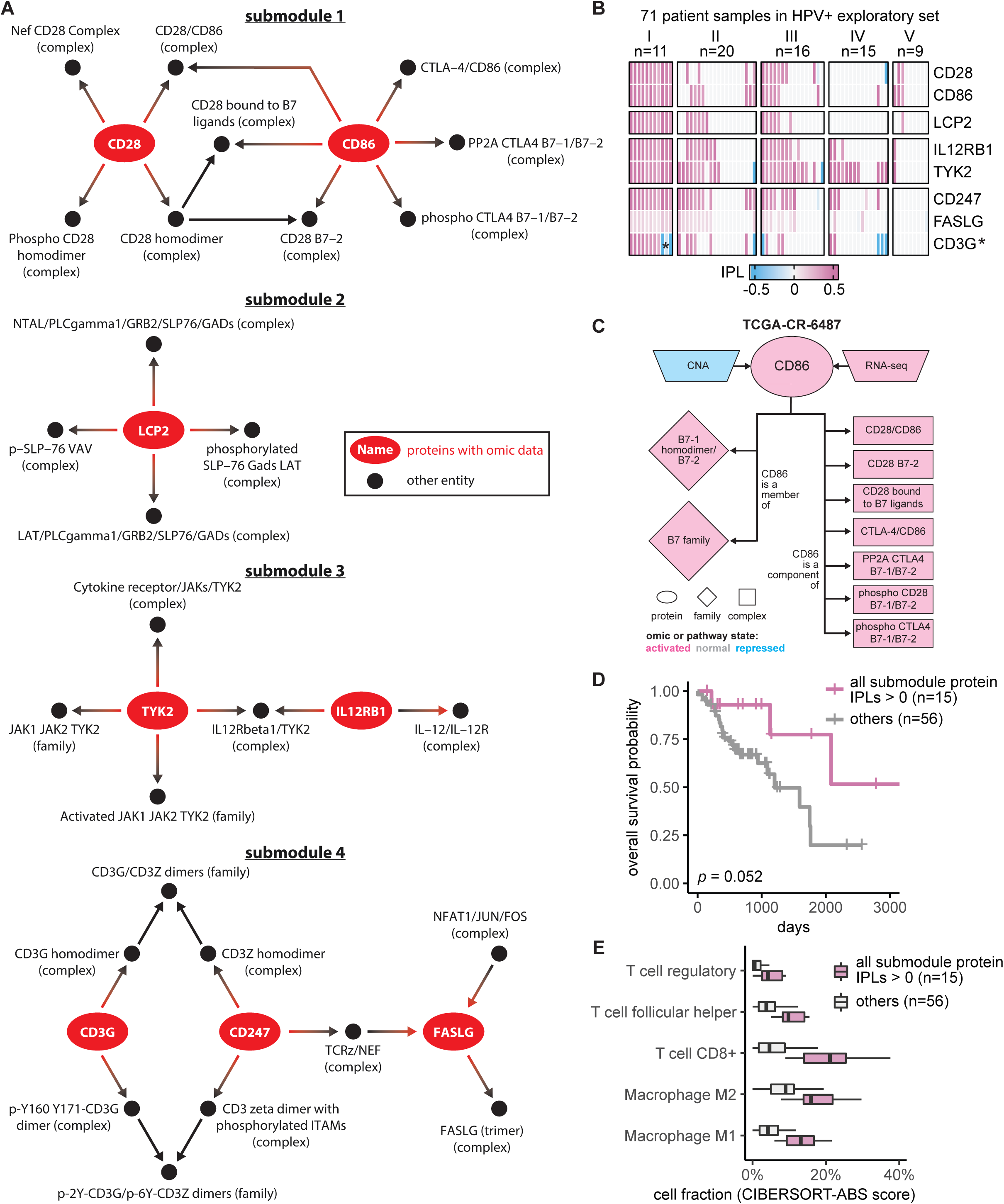
Seven proteins identified in the immune response patient group from HPV+ exploratory set associated with patient overall survival. (**A**) Consensus pathway submodules in the eleven immune response patient samples from group I. Proteins are colored in red and other pathway entities are black; (**B**) inferred pathway levels (IPLs) of submodule proteins in the 71 HPV+ exploratory set samples. Except for CD3G, which had both positive and negative IPLs (denoted by *) in group I, the other seven proteins had positive IPLs; (**C**) CNA, RNA, and pathway states of CD86 as well as pathway states of its pathway network neighbors in a group I patient sample TCGA-CR-6487. B7-1 homodimer/B7-2 (family), B7 family (family), and phospho CD28 B7-1/B7-2 (complex) do not have activated pathway state in all the eleven group I patients (Supplementary Figure 5B) and thus are not included in Figure 3A; (**D**) Overall survival of the HPV+ exploratory set stratified by IPLs of the seven proteins combined; (**E**) HPV+ exploratory set immune cell compositions stratified by IPLs of all seven proteins combined.

To understand what factors determine pathway levels of these seven proteins, we developed an approach for pathway state visualization in MPAC. We transformed the continuous-valued inferred pathway levels to discretized pathway states with the values activated, normal, or repressed. The resulting plots presented a protein’s direct pathway network interaction partners and all associated pathway state information for that protein and its partners, under the reasoning that a determinant of a functionally-implicated protein’s pathway state would have correlated states across all patients (Supplementary Figures 5-7). For example, in all 11 patients, CD86’s pathway states agreed with its RNA states as well as with six of its seven downstream interacting complexes (Supplementary Figure 5B), indicating their parallel roles in determining CD86’s pathway states. In contrast, CD86’s two downstream gene families and one downstream complex had states that disagreed with CD86 in one or three patients, respectively (Supplementary Figure 5B), suggesting a less influential role. CD86’s CNA states, to the other extreme, did not agree with CD86’s pathway state in any patient (Supplementary Figure 5B). In one patient, TCGA-CR-6487, CD86’s CNA state is repressed and its RNA state is activated (Figure 3C). If studying CD86 from individual genomic datasets without any pathway information, it would be hard to determine CD86’s functional protein state, illustrating the advantages of our pathway-based approach. Another feature of MPAC’s visualization function was showing patient-to-patient variations on pathway determinants. FASLG, for instance, had many upstream and downstream neighbors, of which only two upstream and one downstream complexes had pathway states correlated with FASLG, while all the other neighbors had various states across the eleven patients (Supplementary Figures 7B and 8). Such diverse states of FASLG’s neighbors likely reflected subtle cancer mechanism differences within this patient group and MPAC can highlight these differences.

To examine potential clinical implications of these seven proteins, we evaluated their association with the patients’ overall survival. For proteins from the same submodule, we used their inferred pathway levels to divide the 71 HPV+ exploratory patients into two groups: those with all relevant proteins in activated pathway levels and those that were not. This approach facilitates easy visualization to examine the association with survival data, while also taking advantage of the fact that all seven proteins exhibit activated pathway states in Group I patients. The overall survival distributions of patients from the two groups were compared and evaluated by a log-rank test. For every submodule, although the improvement was not always statistically significant, the set of patients with proteins with activated pathway levels always had a better survival distribution than the set that did not (Supplementary Figure 9A). In particular, when patients had both CD247 and FASLG with activated pathway levels, their overall survival was significantly better than those that did not (log-rank *p*=0.00098). Moreover, dividing the same set of patients by the activation of all seven proteins (Figure 3D) or individual proteins (Supplementary Figure 9B) also produced overall survival advantages in all cases but with log-rank *p* ranging from 0.0033 to 0.17. Similar analysis revealed the same trend using the measure of progression-free survival, where patients with the activation of the seven proteins often experience reduced tumor progression, although the association is not as strong as that observed with overall survival in terms of log-rank p (Supplementary Figure 10). Notably, TYK2 and IL12RB1 have been identified by others as associating with HNSCC patient prognosis (He *et al*., 2022; Chen *et al*., 2022). CD28, CD86, CD247, and FASLG have been previously shown to be part of small gene sets that are prognosis-related in HNSCC patients (Zhu *et al*., 2022; Wang *et al*., 2022; Chi *et al*., 2022; de Vos *et al*., 2020). The good association with patient overall survival and reduced tumor progression indicated potential clinical implications of these seven proteins individually and collectively.

Since the seven implicated proteins were identified from the immune response patient group, we explored the relationship between these proteins and immune response. We used a bulk RNA-seq deconvolution method, CIBERSORT in absolute mode (Newman *et al*., 2015), to estimate immune cell composition for the 71 HPV+ exploratory patients and associate them with inferred pathway levels of the seven proteins. CIBERSORT-inferred cell composition was comparable across cell types within the same patient as well as across patients for the same cell type. As in the survival analysis shown in Figure 3D, patients were stratified by whether or not they had all seven proteins with activated pathway levels. For patients in this ‘activated group’, the tumor sample always had substantially higher compositions of T follicular helper cells, CD8+ T cells, regulatory T cells, and M1 and M2 macrophages (Figure 3E; Supplementary Figure 11A). Thus, similar to the prior results cited above, this association indicates that patients with the seven proteins with activated pathway levels usually had higher levels of immune cell infiltration and further suggests that inferred pathway levels of the seven proteins can serve as indicators for immune infiltration.

### Independent validation set confirmed MPAC’s immune response group and key proteins

We used the independent validation set of eighteen HPV+ patients that was held out during MPAC model development and exploratory set analysis to further assess the generality of the seven key proteins identified from the immune response patient group. Thus, we repeated the same MPAC analysis on this validation set, splitting the eighteen validation set patients into two groups. The six patients in the resulting validation group II again had many significantly enriched GO terms related to immune response (Figure 4A). None of the patients with the originally implicated submodule proteins with activated pathway levels died in the interval of record, a notably better overall survival record than those with submodule proteins in normal or repressed pathway levels (Supplementary Figure 12A). Similarly, the overall survival rate of the three patients with all seven proteins in activated pathway levels was always better than the other fifteen patients in the validation set (Figure 4B). The same trend was observed when stratifying patients by individual proteins (Supplementary Figure 12B). The lack of statistically significant differences between two patient groups was due to the small number of patients (Figure 4B; Supplementary Figure 12).

**Figure 4.**
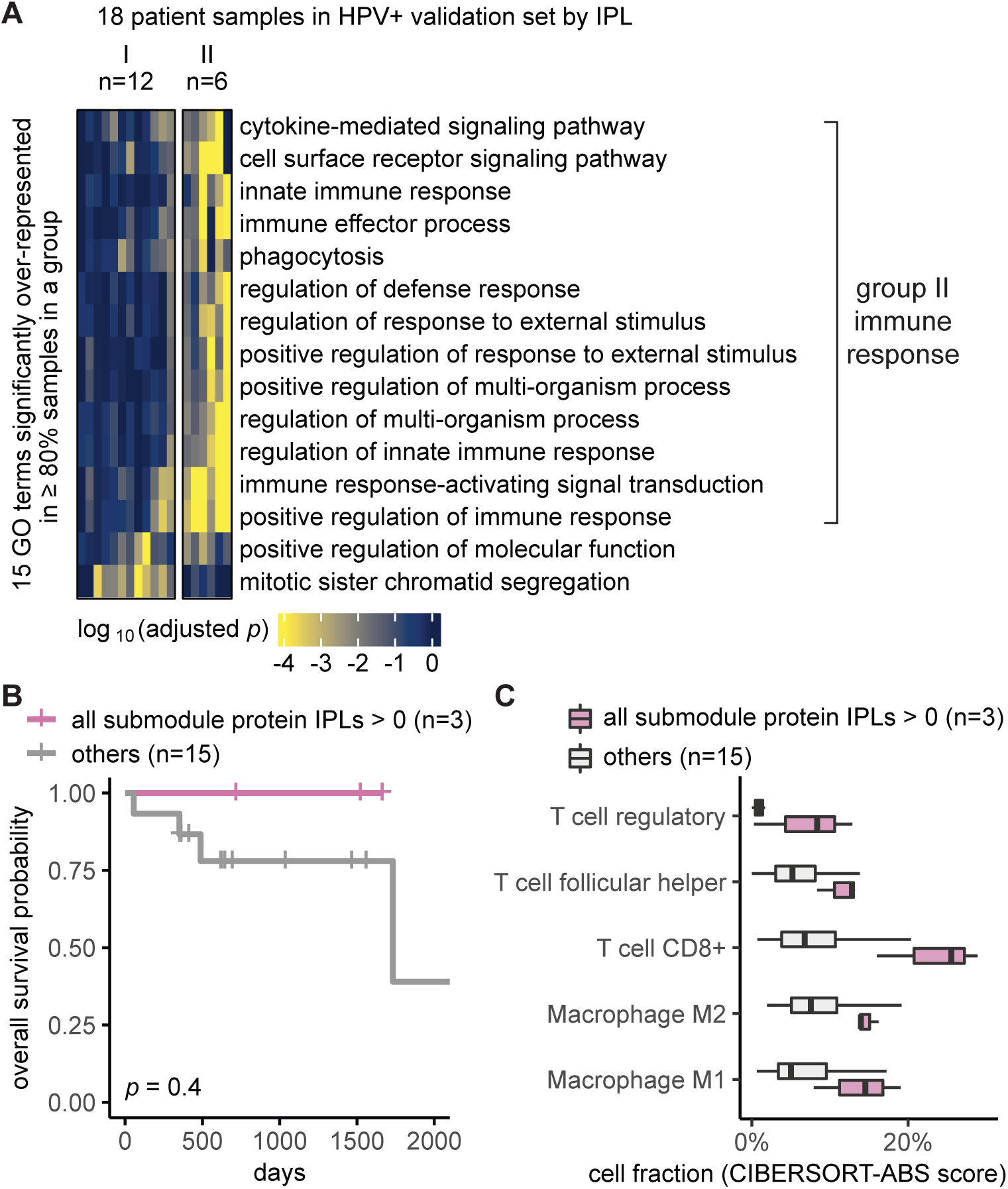
Independent validation set confirmed MPAC’s immune response group and key proteins. **(A)** Grouping of HPV+ validation set patient samples. The selection threshold was lowered to ≥80% in order to include more GO terms; (**B**) Overall survival of the HPV+ validation stratified by the inferred pathway levels (IPLs) of all the seven submodule proteins combined; (**C**) HPV+ validation set immune cell compositions stratified by the IPLs of all the seven proteins combined.

We further examined if the activated pathway levels of the seven proteins are also associated with immune cell infiltration, using the same analysis as for the exploratory set. Validation set patients with the seven proteins in activated pathway levels often had higher fractions of T follicular helper cells, CD8+ T cells, regulatory T cells, M1 and M2 macrophages (Figure 4C), the same as we observed in the exploratory set, although the difference was not significant most likely due to the small sample size (Supplementary Figure 11B). Altogether, the independent validation set supported MPAC’s predictions in the exploratory set and greatly reduced the possibility of bias from using the exploratory set alone.

### MPAC’s advantages over PARADIGM and robustness

To assess MPAC’s benefits over PARADIGM, we asked whether PARADIGM can also discover an immune response patient group as MPAC does. We downloaded PARADIGM’s inferred pathway levels from NCI’s PanCanAtlas website. 70 of the 71 patients in the HPV+ exploratory set and all 18 patients in the validation set have PARADIGM inferred pathway levels available. We applied Gene Set Enrichment Analysis (GSEA) on the same set of GO terms used by MPAC and then clustered patients by K-means based on the GSEA results. For the HPV+ exploratory set, we tried for two to five groups, and two groups appeared to be a good choice because it does not have any group with just one or two samples (Supplementary Figure 13A). Only two GO terms are significantly overrepresented in ≥ 50% of samples in Group c2 and none for Group c1 (Supplementary Figure 13B). The two GO terms are related to development, which is similar to Group V by MPAC (Figure 2). PARADIGM does not find an immune response group. For the HPV+ validation set, we also tried two to five groups, and two groups were taken for the same reason as for the exploratory set (Supplementary Figure 13C). Six GO terms are significantly overrepresented in ≥ 80% of samples of Group c2 and none for Group c1 (Supplementary Figure 13D). All the six GO terms are related to development of morphogenesis. Once again, PARADIGM does not find an immune response group. In summary, MPAC shows advantages over PARADIGM because MPAC recovers a unique immune response patient group and PARADIGM’s largest clusters in both the HPV+ exploratory and validation sets are not enriched for any GO terms.

MPAC is robust to many of the analysis configuration choices. First, we did a subsampling analysis by randomly taking 10%, 30%, and 50% of the 89 HPV+ exploratory and validation sets. We applied MPAC on each of them and found that an immune response patient group can be obtained at both 30% and 50%, but not at 10% (Supplementary Figure 14), indicating MPAC may not generate consistent results for sample size < 10.

Secondly, we created multiple data splits that swapped which of the 89 HPV+ samples were used as the exploratory and validation set, similar to 5-fold cross-validation in supervised learning. The existing HPV+ exploratory (Figure 2) and validation (Figure 4) sets were used as split #1. The same MPAC protocol and parameters were applied to splits #2 to #5, so there is no supervised learning per se on the exploratory sets from these four splits. Most of the results generated an immune response group except for split #2’s validation set, split #5’s exploratory and validation sets (Supplementary Figure 15). Split #3’s validation set has Group c2 as an immune response group and this group also contains significantly overrepresented cell cycle GO terms (Supplementary Figure 15D). Overall, this exploratory and validation set resampling result demonstrated the robustness of MPAC.

Thirdly, we examined how changing MPAC settings affects the results. The default 100 permutations are needed when the sample size is small as the 18 patients in the HPV- validation set, but it can be reduced to 50 or 20 for a larger sample size like the 71 patients in the exploratory set (Supplementary Note 2; Supplementary Figure 16). The default two standard deviation threshold to define input RNA states can be increased to three, but decreasing to one resulted in largely different results (Supplementary Note 3; Supplementary Figure 17). Including more patient-specific pathway sub-networks in MPAC’s Step 5 is unlikely to affect the results, because the second largest sub-networks are much smaller than the largest one (Supplementary Note 4; Supplementary Figure 18). Moreover, such an approach does not bias the output toward ubiquitously overrepresented GO terms (Figure 2 and 4A). Integrating CNA with RNA-seq data boosted MPAC, because CNA has a stronger impact than RNA-seq on determining a protein’s pathway state overall (Supplementary Note 5; Supplementary Figure 19). Separating HPV+ and HPV- patients before applying MPAC is recommended when the input pathway file has little information on HPV-specific pathways (Supplementary Note 6; Supplementary Figure 20).

Lastly, we applied MPAC to a different TCGA cancer type to evaluate its generalizability. We chose cholangiocarcinoma because the original study on this cohort (Farshidfar *et al*., 2017) reported pathway analysis results only from bulk RNA-seq data, not multi-omic data. MPAC’s result on the 35 cholangiocarcinoma samples that have both CNA and RNA-seq data shows three groups with distinct biological functions (Supplementary Figure 21). Group c1 is mainly on metabolic processes, especially on xenobiotic metabolic processes. Group c2 is mainly on apoptotic process and response to unfolded protein. Group c3’s function is unclear because it does not have any GO term significantly overrepresented in ≥ 80% of samples (i.e. ≥ 8 samples). These three groups illustrate MPAC’s applicability to a cancer type other than HNSCC. The significantly overrepresented GO terms in this cohort are different from those in HNSCC, largely because of different disease mechanisms. Additionally, 74 and 53 submodule proteins were identified in Group c1 and c2, respectively. Because the number of submodule proteins exceeds the small sample size of 35, we did not pursue survival analysis in this cohort.

### An interactive MPAC Shiny app supports visualization of results and new analyses

We built an R Shiny app (https://github.com/pliu55/MPAC_Shiny) to display all the results generated from this work and support new analyses of the data. It shows enrichment results from 2,805 pathways, inferred pathway levels of 19,477 pathway entities, CNA and RNA states of 6,251 pathway proteins, and overall survival and immune cell compositions of 492 HNSCC patient samples. Moreover, it illustrates a protein’s pathway membership and network neighbors. On the landing page’s sidebar, users can choose one of the four TCGA-HNSCC datasets: HPV+ or HPV- combined with an exploratory or validation set. The MPAC app presents results at both the pathway- and protein-level. On the pathway-level page (Figure 5A), Shiny app Box (i) displays pathway enrichment results similar to the ones shown in Figure 2, Figure 4A, and Supplementary Figure 1A. Users can enter any pathway(s) of interest to look at their enrichments in MPAC-defined patient groups. To understand which proteins lead to a pathway enrichment, Box (ii) shows inferred pathway levels of all the proteins from a pathway. For example, in the pathway ‘positive regulation of interleukin-2 biosynthetic process’, CD28, CD3E, CD4, CD80, CD86, and PTPRC have positive inferred pathway levels in a majority of group I HPV+ exploratory patient samples (Figure 5A), suggesting they are the determinants resulting in this pathway’s enrichment in group I patients.

**Figure 5.**
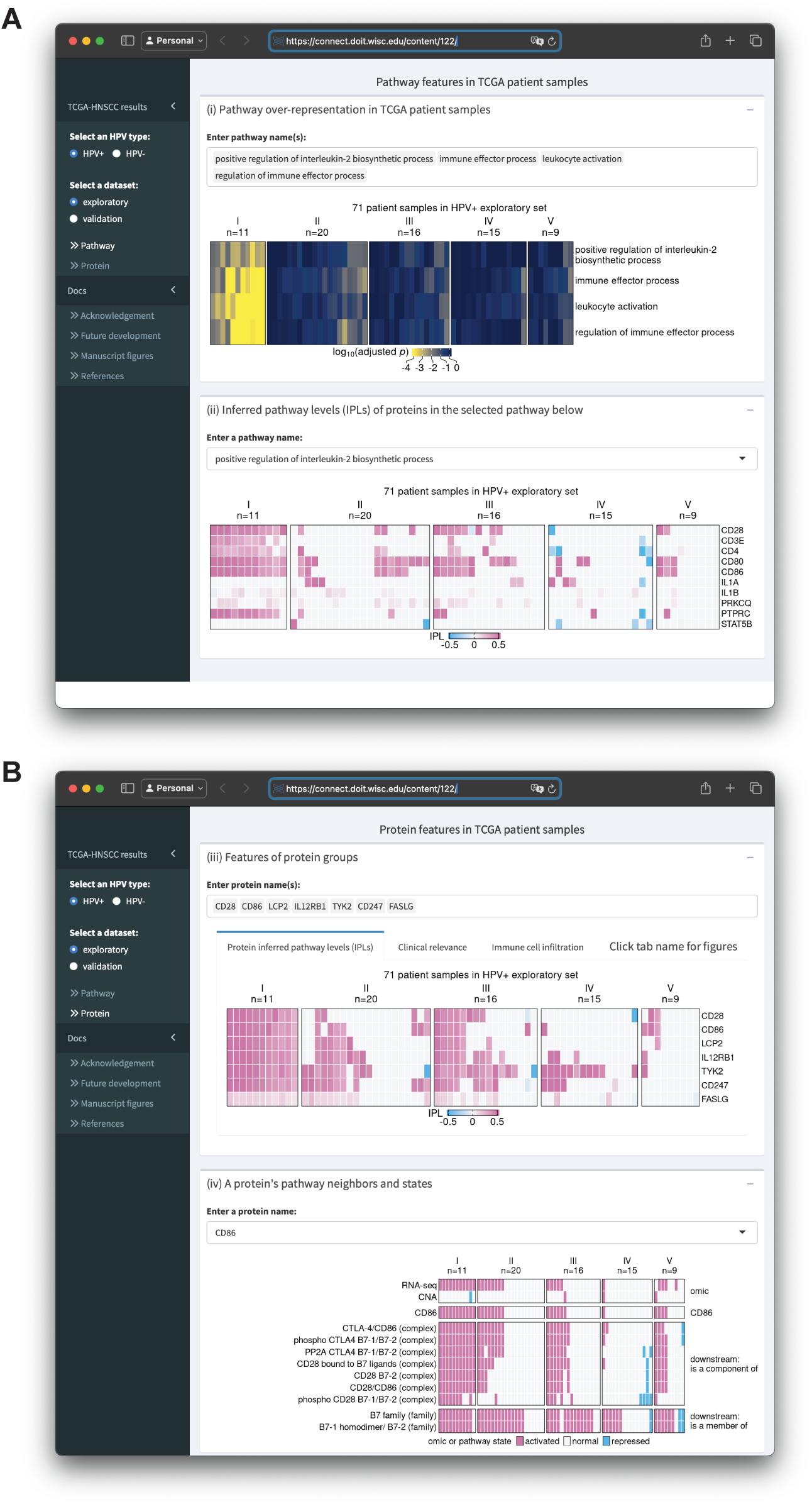
Screenshots of an R Shiny app displaying MPAC results from the HPV+ exploratory set. (**A**)The upper box shows enrichment of multiple user-selected pathways and the lower box shows protein inferred pathway levels (IPLs) from a user-selected pathway; (**B**) The upper box shows IPLs of multiple user-selected proteins and the lower box shows the pathway states of a user-selected protein and its pathway neighbors as well as its CNA and RNA state.

In Box (ii), users can enter or select any pathway of interest to examine their proteins’ inferred pathway levels. At the protein-level page (Figure 5B), Box (iii) contains results for a group of user-specified proteins. It has three tabs displaying proteins’ inferred pathway levels, overall survival and immune cell composition of patients stratified by proteins’ inferred pathway levels. These figures are similar to Figures 3B, 3D, 3E, 4B, and 4C; Supplementary Figures 9, 11, and 12, with the flexibility of showing the results for any user-specified protein(s) on any of the four TCGA-HNSCC datasets. Box (iv) shows a heatmap of the CNA, RNA, and pathway states for any user-entered protein as well as pathway states of this protein’s pathway network neighbors. It is similar to Supplementary Figures 5-7 with the same flexibility as Box (iii). In addition to the interactive data display, this app also contains documentation regarding re-generating figures presented in this manuscript, related papers, future developments, and acknowledgement. In summary, the MPAC Shiny app provides a convenient way to explore all the results generated from this work, especially those not presented as figures in this manuscript.

## Discussion

We presented MPAC as a computational framework with several unique features compared to other pathway-based multi-omic integration tools (Supplementary Table 3). MPAC calculates inferred pathway levels, predicts patient groups with biologically meaningful pathway profiles, and identifies key proteins with potential clinical associations. One group of HNSCC patients was predicted to have alterations in immune response pathways. This group could not be identified from CNA or RNA-seq data alone or by PARADIGM. This finding illustrates the advantages of our pathway-based multi-omic approach. MPAC can use prior knowledge of pathway interactions in the form of a pathway network to integrate CNA and RNA-seq data and infer proteins’ pathway behavior. A protein’s pathway behavior from MPAC was not solely inferred from its CNA or RNA but also its pathway neighbors (Figure 3C; Supplementary Figures 5-8). Analysis based on CNA and RNA-seq data alone would miss this important biological principle.

Our analysis showed that MPAC can predict patient groups with potentially relevant clinical properties by their pathway profiles. The results presented above (Figure 2 and Supplementary Figure 1) identified an immune response patient group in both HPV+ and HPV- HNSCC, which echoes a new subtype defined by recent studies. One study (Huang *et al*., 2021) applied a proteogenomic approach on 108 HPV- HNSCC patients. By considering CNA, RNA, miRNA, protein, and phosphopeptide data, the authors defined three subtypes of HNSCC: chromosome instability, basal, and immune. These subtypes mirrored the immune response patient groups MPAC identified in both HPV+ and HPV- patients. Moreover, the authors analyzed immune-hot tumors and revealed the presence of both cytotoxic immune cells (e.g., CD8+ T cells and M1 macrophages) and immunosuppressive cells (e.g., regulatory T cells and M2 macrophages). This is consistent with our analysis of HPV+ tumors stratified by inferred pathway levels of the seven proteins identified from the immune response patient group. Tumor samples with activated pathway levels of any of the seven proteins always had a higher fraction of CD8+ T cells, regulatory T cells, and M1 and M2 macrophages in both the exploratory and validation sets (Figures 3E and 4C). Second, a multi-omic analysis of thirty-three TCGA cancer types (Tiong *et al*., 2022) identified gene groups enriched by immune response as well as cell cycle, which were also observed in MPAC’s results (Figures 2 and 4A; Supplementary Figure 1A). The agreement between these two studies supports MPAC’s discovery of the immune response patient group.

Further, not only did the seven proteins identified by MPAC associate with immune cell composition, but their activated pathway levels also associated with better overall survival. This was demonstrated in our exploratory patient set (Figure 3D and Supplementary Figure 9) and supported by our validation set (Figure 4B and Supplementary Figure 12). The corroboration by the validation set illustrates a major strength of MPAC. To understand these seven proteins’ clinical values and whether they could serve as biomarkers would require a prospective patient cohort, which is not available to us currently (As of May 6, 2024, according to cBioportal (de Bruijn *et al*., 2023), https://www.cbioportal.org/), the only HNSCC dataset that has both CNA and RNA-seq data available is from TCGA, which is used in this work). However, the analyses here demonstrated how the MPAC software could be applied in a prospective setting.

MPAC has several advantages over the PARADIGM algorithm that it calls as a subroutine. When preparing the input for predicting pathway levels, MPAC uses a data-driven approach to define each gene’s discrete states based on both tumor and normal tissue samples, whereas PARADIGM arbitrarily assigns the top, middle and lower third of omic-ranked genes as activated, normal and repressed (Vaske *et al*., 2010). MPAC also provides downstream analyses on inferred pathway levels, including built-in permutation testing, defining altered pathways, predicting patient groups, and identifying key proteins with potential clinical implications. All of these functions have been implemented in an R package available through Bioconductor making it easier for others to use in their studies. The MPAC R Shiny app also supports convenient visualizations of the MPAC predictions.

Multi-omic integration methods have been developed for diverse applications (Maghsoudi *et al*., 2022; Zitnik *et al*., 2024), such as embedding single-cell data (Ashuach *et al*., 2023; Argelaguet *et al*., 2020), clustering cancer samples (Chauvel *et al*., 2020; Wang *et al*., 2014), and pathway reconstruction (Tuncbag *et al*., 2016; Winkler *et al*., 2022; Paull *et al*., 2013). Multi-omics analyses have been particularly prominent in cancer, with pathway enrichment (Paczkowska *et al*., 2020), representation learning (Leng *et al*., 2022), supervised prediction of cancer subtypes or patient outcomes (Poirion *et al*., 2021; Choi and Chae, 2023), and biologically interpretable neural networks (Wysocka *et al*., 2023) as representative areas of study. MPAC’s unique role in this methodological landscape is that through PARADIGM it directly uses pathway interactions to combine information across omic data types, learn protein activities, and conduct downstream analysis with those protein activities.

In this work, we limited the input multi-omic data to CNA and RNA-seq, given PARADIGM’s previous success with these two data types. With the availability of many other types of omic data from TCGA and the Clinical Proteomic Tumor Analysis Consortium (Huang *et al*., 2021) on large cohorts of cancer patients, time course multi-omic data (G. Stein-O’Brien *et al*., 2018), single-cell RNA-seq (Puram *et al*., 2017), spatial transcriptomics (Li *et al*., 2024; Lee *et al*., 2024), and spatial proteomics (Causer *et al*., 2023), one of our future goals is to make MPAC compatible with as many omic data types as possible. This requires extending the MPAC software as well as the input biological pathways to include knowledge on the relevant molecules and associated regulatory mechanisms. Expanding the input biological pathways will also help disease-specific studies as shown in our analysis on separating HPV+ and HPV- HNSCC patient samples (Supplementary Note 6; Supplementary 20). For studies focusing on a specific disease or condition, smaller and pertinent input pathways will expedite MPAC’s PARADIGM subroutine calculations on permuted data (Supplementary Note 7; Supplementary Figure 22).

## Methods

### Genomic and clinical datasets

We downloaded the TCGA HNSCC genomic datasets (Cancer Genome Atlas Network, 2015) from NCI GDC Data Portal version 29.0 (https://portal.gdc.cancer.gov/), which was released on March 31, 2021. Gene-level copy number scores were used for CNA and log_10_(FPKM+1) values were used for RNA-seq. Patients’ HPV status was obtained from their biospecimen manifest files. Patients’ clinical data was downloaded from TCGA Pan-Cancer Atlas (Liu *et al*., 2018) via https://api.gdc.cancer.gov/data/1b5f413e-a8d1-4d10-92eb-7c4ae739ed81. 492 HNSCC patients that had CNA, RNA-seq, and clinical data available were stratified by HPV status and then randomly divided into exploratory sets (71 HPV+ and 322 HPV-) and validation sets (18 HPV+ and 81 HPV-). Importantly, only the exploratory set was used for all MPAC algorithm development and refinement.

MPAC’s pathway definitions were taken from the TCGA Pan-Cancer Atlas (Hoadley *et al*., 2018), which compiled interactions from NCI-PID (Schaefer *et al*., 2009), Reactome (Gillespie *et al*., 2022), and KEGG (Kanehisa *et al*., 2023) and superimposed them into a single network. The input network for MPAC included 19,477 entities, including 7,321 proteins, 9,349 complexes, 2,092 families, 591 abstract processes, 15 miRNAs, 82 RNAs, and 27 other types of entities. It also included 45,313 interactions containing 2,133 activations and 401 repressions at the transcript-level, 7,723 activations and 1,083 repressions at the protein-level, 24,870 and 9,103 memberships for complexes and families, respectively. The 2,085 BIological Process GO terms (Gene Ontology Consortium *et al*., 2023) for characterizing patient or cell line pathway alteration were downloaded from the DrugCell (Kuenzi *et al*., 2020) GitHub repository (https://github.com/idekerlab/DrugCell/blob/public/data/drugcell_ont.txt). GO terms from DrugCell had more distinct genes between parental and offspring terms because it required a parent to have ≥ 10 genes distinct from all child terms and have ≥ 30 genes more than any child. The root GO term (i.e., the ancestor of all the other GO terms), ‘biological process’, was not used in this study because it was not a specific functional description.

### MPAC workflow

For TCGA data, the signs of CNA focal scores were used to define activated, normal (i.e., focal score is exactly zero), or repressed CNA state as the input for MPAC. To define input RNA state, a gene’s RNA-seq expression levels from normal patient samples were fit to a Gaussian distribution. If a gene’s expression levels in tumor samples fell within two standard deviations from the mean of this distribution, the gene’s RNA state was defined as normal. Otherwise, its RNA state was repressed or activated depending on whether its expression level was below or above the two standard deviations from the mean. MPAC takes two standard deviations on a Gaussian distribution as a threshold because it corresponds to the common p < 0.05 cutoff.

MPAC ran PARADIGM in the default configuration (Vaske *et al*., 2010) except with a more stringent expectation-maximization convergence criteria of change of likelihood < 10^-9^ under a maximum of 10^4^ iterations. To prepare permuted input, paired CNA and RNA states were randomly shuffled between all the genes within the patient. 100 permuted samples were prepared per each real tumor sample resulting in 49,200 permuted samples in total for the 492 patients. This large number of computational jobs were processed through UW-Madison’s Center for High Throughput Computing (Center for High Throughput Computing, 2006) with HTCondor (Thain *et al*., 2005).

A pathway entity’s inferred pathway level from a real tumor sample was set to NA if it fell within three median absolute deviations of the inferred pathway levels from the corresponding 100 permuted samples. This filtering helped to remove inferred pathway levels that could be observed by chance. Entities with non-NA inferred pathway levels were mapped to the input pathway network. The largest connected subset of the pathway network with non-NA inferred pathway levels was kept for downstream analysis. Other entities not in this largest subset had their inferred pathway levels set to zero. This allowed us to focus on the entities that act together in pathways.

After the filtering by permuted samples and the largest pathway subset, an entity’s pathway state was defined by the sign of its inferred pathway level, where a positive or negative inferred pathway level corresponded to an activated or repressed state, respectively, and a zero inferred pathway level corresponded to a normal state. Based on normal or altered pathway states, GO enrichment was calculated by Fisher’s exact test, and the p-values were adjusted by the Benjamini and Hochberg procedure. Similarly, GO enrichment was calculated for the CNA and RNA inputs by their normal or altered states. If other gene sets are preferred instead of GO terms, users can supply a GMT-formated file of the gene sets to the ‘fgmt’ option in MPAC’s ‘ovrGMT()’ function.

Patients were grouped by their adjusted *p*-values from GO enrichment based on CNA, RNA, or inferred pathway levels. A clustering method originally designed for single-cell RNA-seq analysis was adapted, where a patient tumor sample was treated as a cell and the |log_10_(adjusted *p*)| was treated as a gene’s expression level. Gene variance was modeled by the modelGeneVar function from the scran package (Lun *et al*., 2016) (version 1.20.1), and the top 100 genes were selected. Patients were grouped by the Louvain method from the igraph R package (Csárdi *et al*., 2024) (version 1.2.11) with 10 or 20 nearest neighbors for HPV+ or HPV-, respectively. Patient groups do not necessarily have similar sizes (Figure 2; Figure 4A; Supplementary Figure 21). Changes in the igraph R package starting with version 1.3 affected the reproducibility of our results but not the main conclusions from our analyses (Supplementary Note 8). The most time-consuming part of MPAC is running PARADIGM using the Pan-Cancer Atlas pathways on real and permuted data. It took a maximum of 4 days to run on real data and two weeks on permuted data using 500∼600 MB memory on each sample (Supplementary Note 7; Supplementary Figure 22).

To summarize pathway features for a group, we plotted heatmaps of log_10_(adjusted *p*) values for GO terms with adjusted *p* < 0.05 in 100% of the patients from the same group (e.g., Figure 2 and Supplementary Figure 1A). When very few GO terms met this criterion, we lowered the percentage threshold (e.g., ≥ 80% in Figure 4A, ≥ 60% in Supplementary Figure 1C) or by specifying a minimum number (e.g., ≥ 3 in Supplementary Figure 1B) of patients in order to include more GO terms.

MPAC offers several functions for users to explore the networks generated during the analysis. The largest network subset can be obtained by MPAC’s ‘subNtw()’ function, while consensus pathway submodules can be obtained by MPAC’s ‘conMtf()’ function and visualized by MPAC’s ‘pltConMtf()’ function.

For survival analysis, we used the inferred pathway levels of one or multiple proteins to stratify patients into two groups: those with all the protein(s) in activated pathway states (i.e., positive inferred pathway level values) and those not. A log-rank test *p*-value was calculated to compare the survival distribution of the two groups.

## Software and data availability

The MPAC package is available at Bioconductor (https://bioconductor.org/packages/MPAC) and archived on Zenodo (https://doi.org/10.5281/zenodo.10805479). MPAC’s Shiny app is accessible at https://connect.doit.wisc.edu/content/122/. The source code for MPAC’s Shiny app is available at GitHub (https://github.com/pliu55/MPAC_Shiny) and archived on Zenodo (https://doi.org/10.5281/zenodo.11623974). Both the R package and Shiny app are available under the GPL-3.0 license.

## Acknowledgements

We acknowledge funding from University of Wisconsin Carbone Cancer Center Support Grant P30 CA014520, from the UW Institute for Clinical and Translational Research Pilot Award UL1 TR002373 from NIH/NCATS, and from a joint pilot program of the Morgridge Institute for Research and the NIH/NCI UW Comprehensive Cancer Center Support Grant; A.G. and I.M.O. acknowledge pilot awards from the Wisconsin Head and Neck Cancer SPORE Career Enhancement Program P50 DE026787; I.M.O. and P.L. acknowledge support from the Wisconsin Head and Neck Cancer SPORE as part of the Biostatistics and Bioinformatics Core P50 CA278595; I.M.O. acknowledges support from the UW Institute for Clinical and Translational Research KL2 TR002374 from NIH/NCATS and the NIH/NCI grant P01 CA250972; P.A. acknowledges support from the NIH/NCI grant P01 CA022443. This research was partially performed using the compute resources and assistance of the UW–Madison Center for High Throughput Computing in the Department of Computer Sciences. A.G. and P.A. are investigators of the Morgridge Institute for Research and an investigator and the director, respectively, of the John and Jeanne Rowe Center for Research in Virology and gratefully acknowledge their support. We thank H. Adam Steinberg for creative contributions to figure design and data visualization, Alicia Williams for organization feedback and manuscript editing, and David Merrell, Deric Wheeler, Mari Iida, Paul Lambert, and Randall Kimple for project feedback.

## Supplementary Materials

### Supplementary figure legends

**Supplementary Figure 1.**
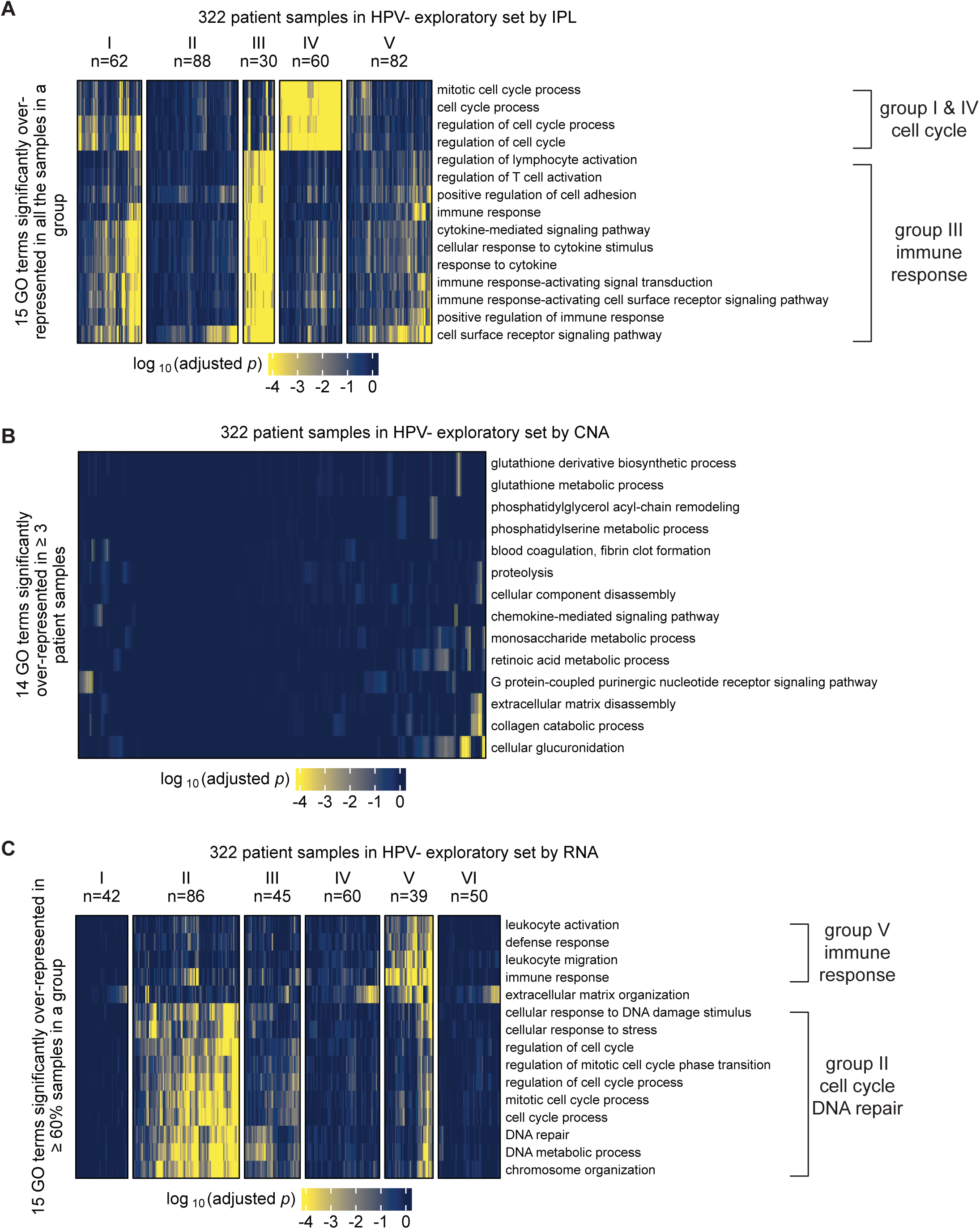
HPV- exploratory set patient samples grouped by GO term enrichment based on IPL (A), CNA (B), or RNA (C). The selection threshold was lowered to ≥ 3 (**B**) or ≥ 60% (**C**) to include more GO terms to avoid bias.

**Supplementary Figure 2.**
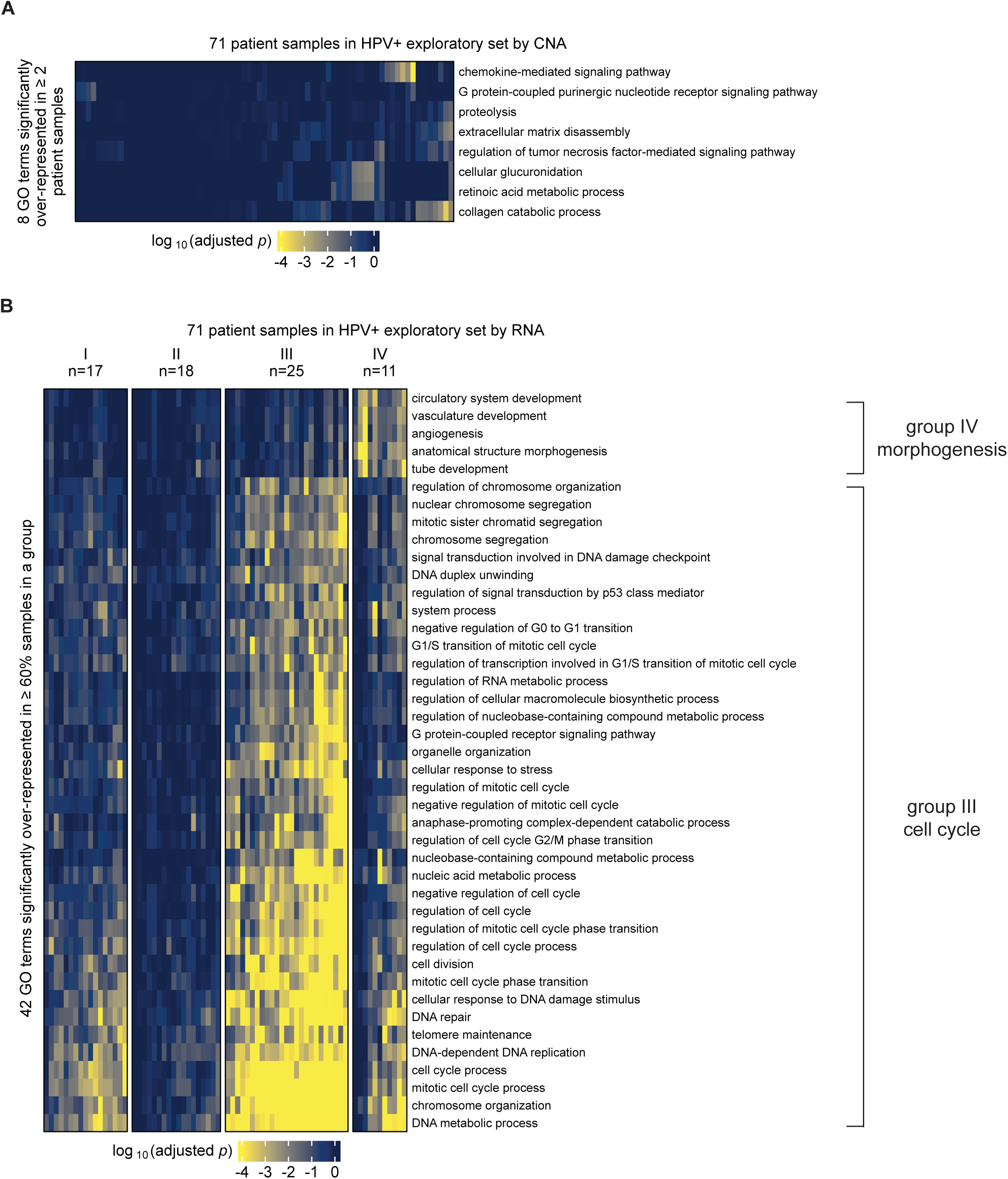
HPV+ exploratory set patient samples grouped by GO term enrichment based on CNA (A) or RNA (B). The selection threshold was lowered to ≥ 2 (**A**) or ≥ 60% (**B**) to include more GO terms to avoid bias.

**Supplementary Figure 3.**
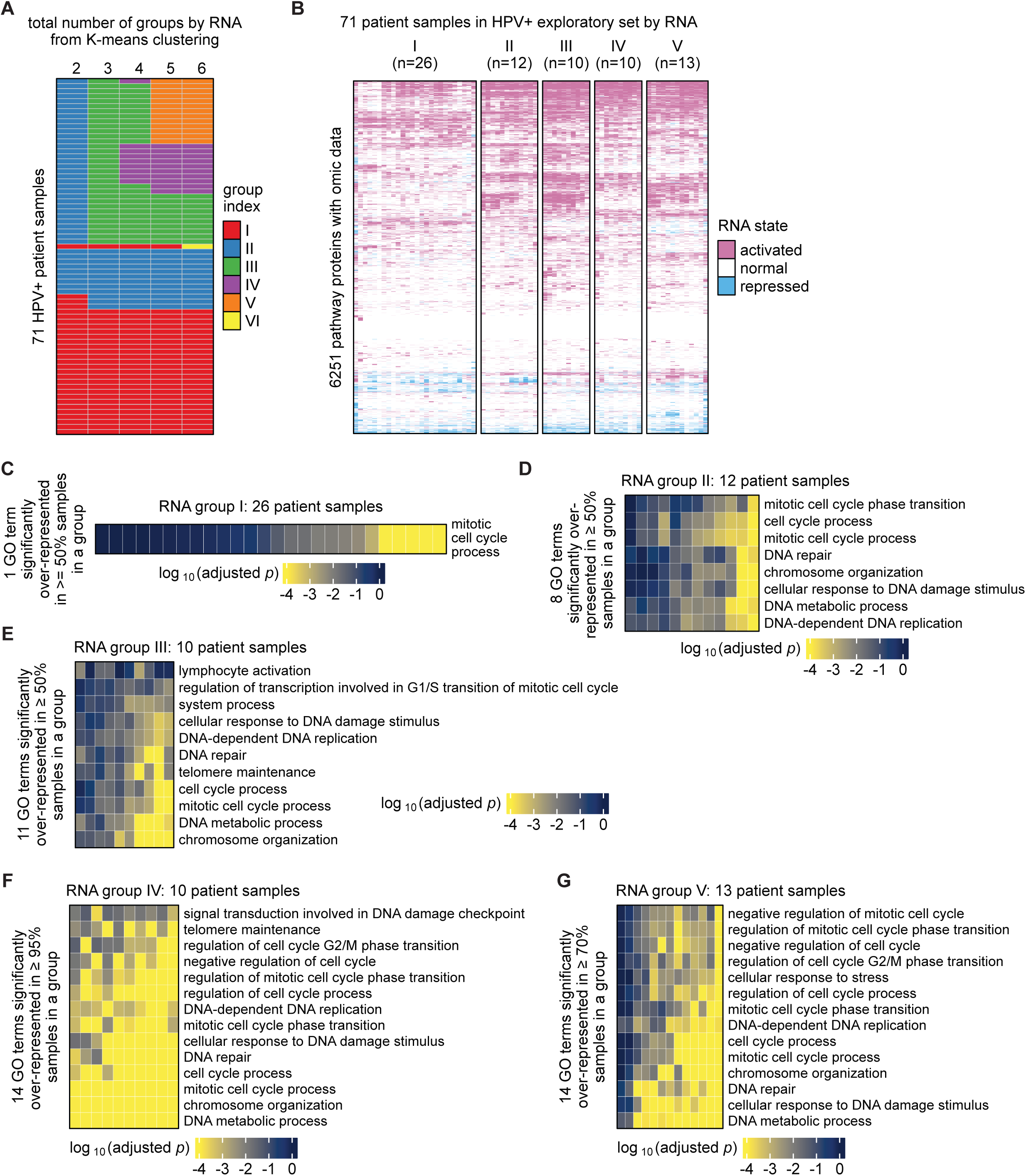
HPV+ exploratory set patient samples grouped by MPAC’s input RNA states. **(A)** Clustering results by K-means under a pre-specified total number of groups ranging from two to six; (**B**) Clustering results by K-means with a pre-specified five groups; (**C**– **G**) Top significantly enriched GO terms in group I (**C**), II (**D**), III (**E**), IV (**F**), and V (**G**). The selection threshold was lowered to ≥ 95% (**F**), ≥70% (**G**), or ≥50% (**C–E**) in order to include more GO terms to avoid bias.

**Supplementary Figure 4.**
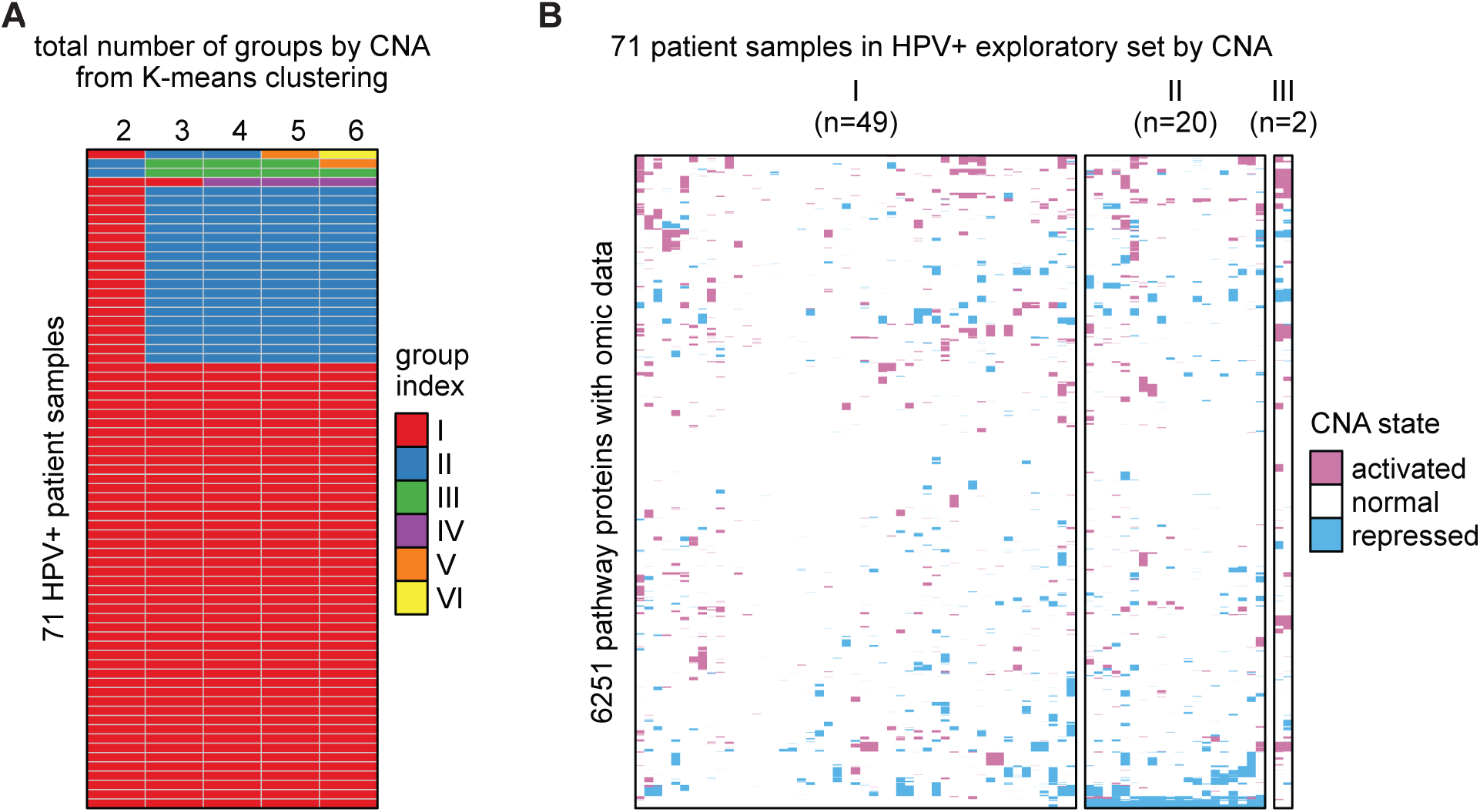
HPV+ exploratory set patient samples grouped by MPAC’s input CNA states. **(A)** Clustering results by K-means under a pre-specified total number of groups ranging from two to six; (**B**) Clustering results by K-means with a pre-specified three groups.

**Supplementary Figure 5.**
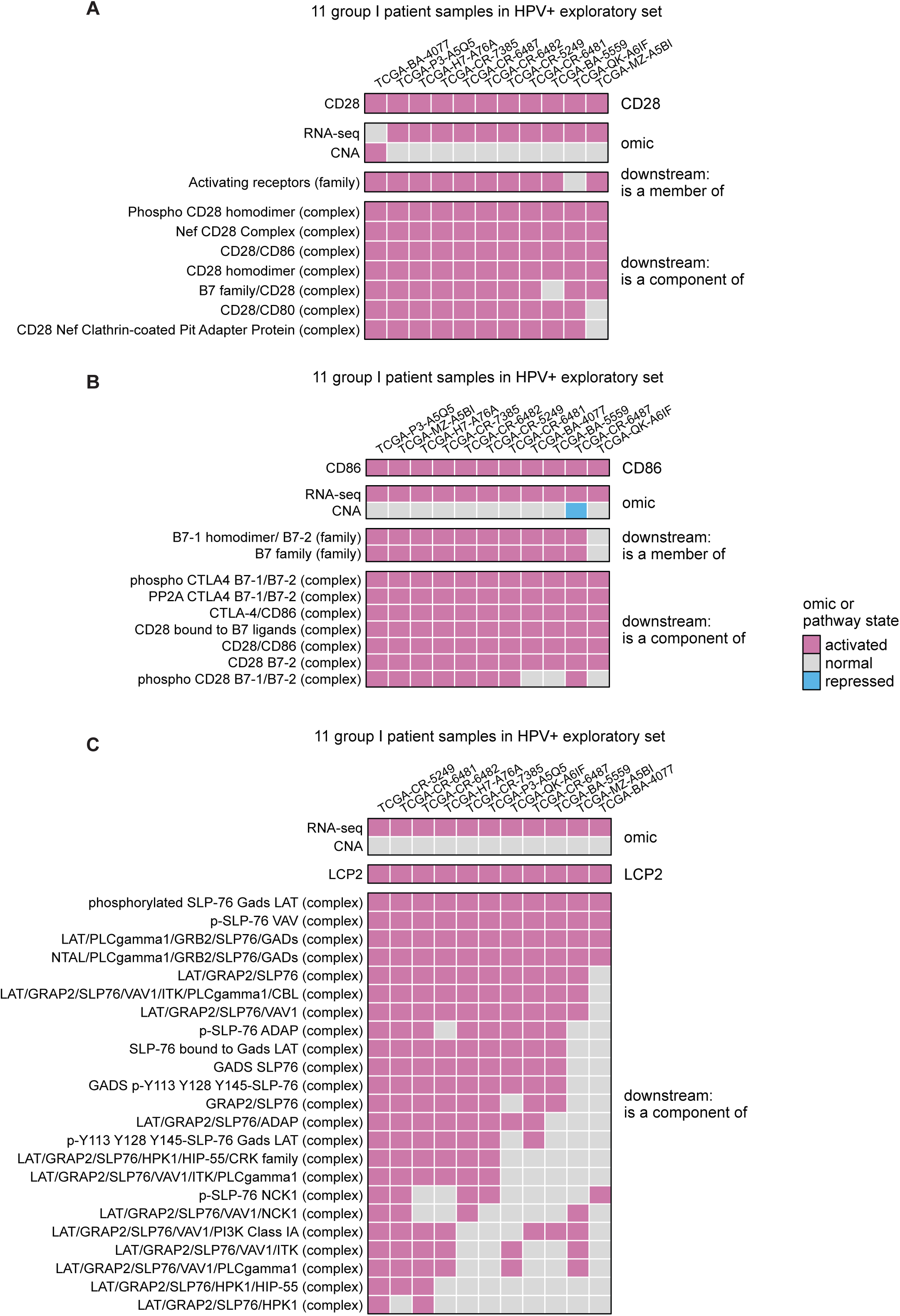
CNA, RNA, and pathway states of CD28 (A), CD86 (B), and LCP2 (C), as well as pathway states of their pathway network neighbors in the eleven group I patients.

**Supplementary Figure 6.**
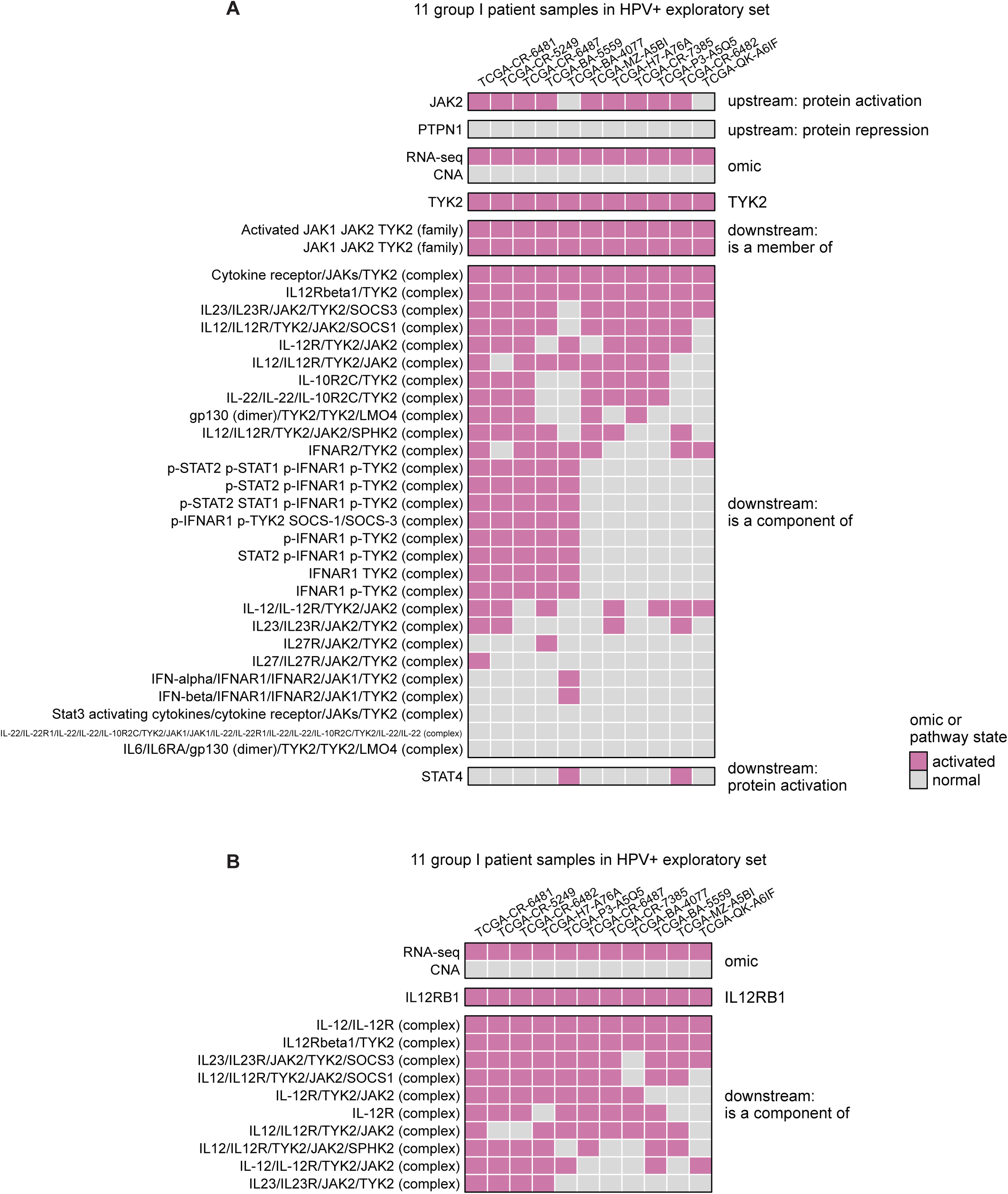
CNA, RNA, and pathway states of TYK2 (A) and IL12RB1 (B), as well as pathway states of their pathway network neighbors in the eleven group I patients.

**Supplementary Figure 7.**
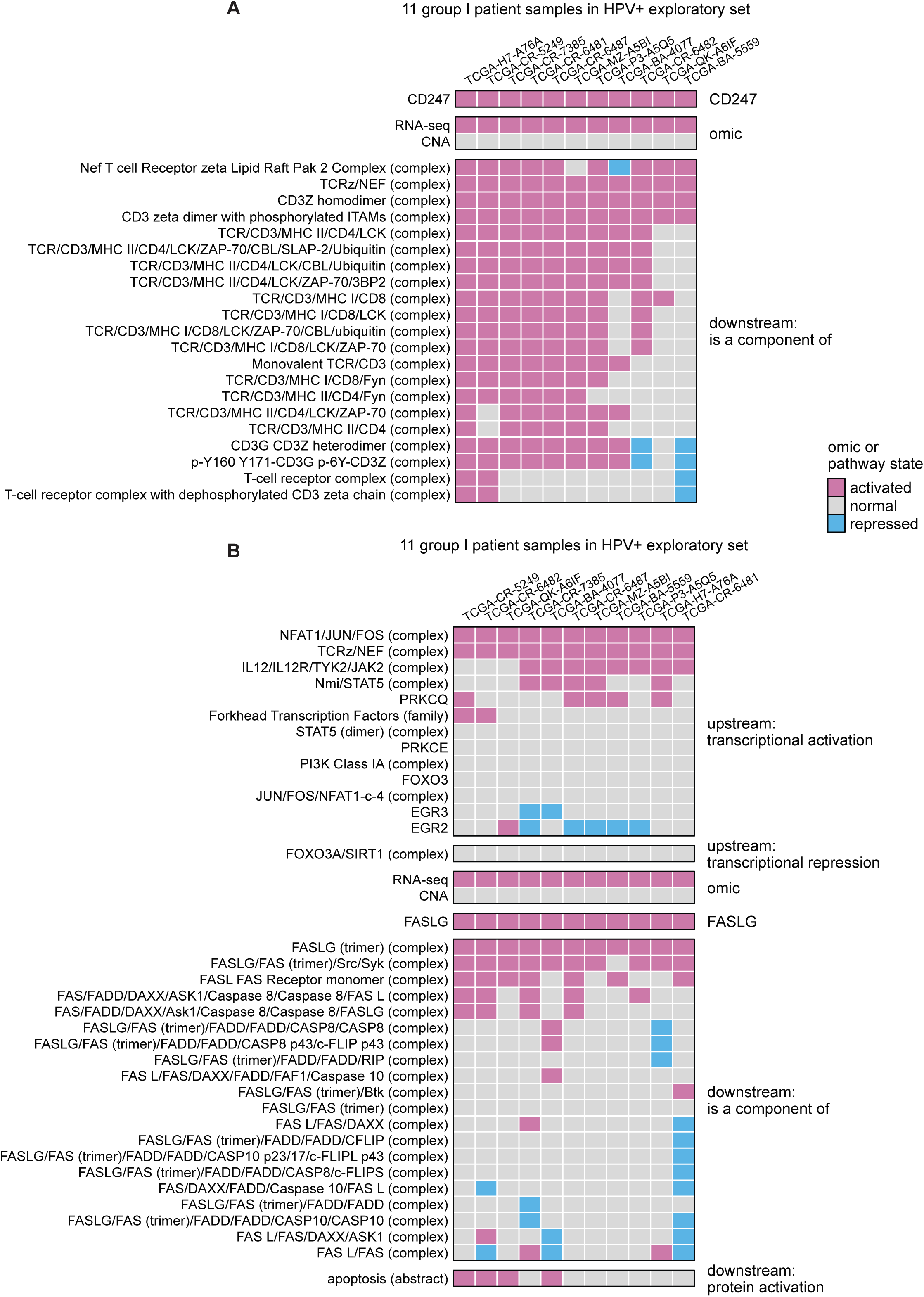
CNA, RNA, and pathway states of CD247 (A) and FASLG (B), as well as pathway states of their pathway network neighbors in the eleven group I patients.

**Supplementary Figure 8.**
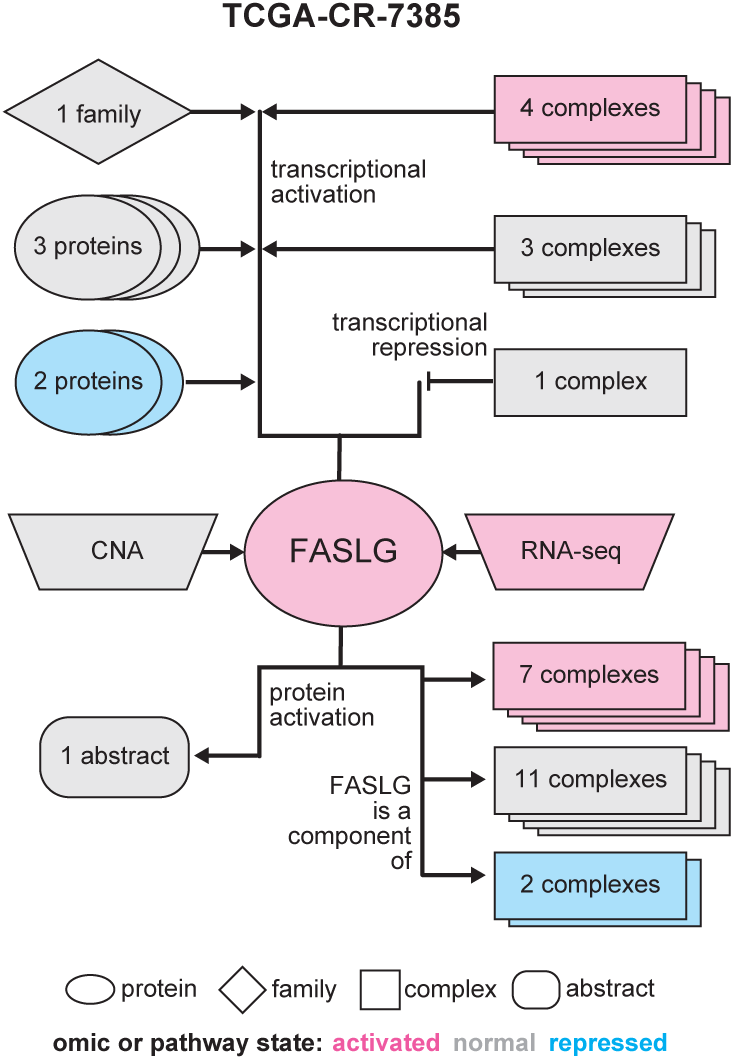
CNA, RNA, and pathway states of FASLG as well as pathway states of its pathway network neighbors in a group I patient TCGA-CR-7385.

**Supplementary Figure 9.**
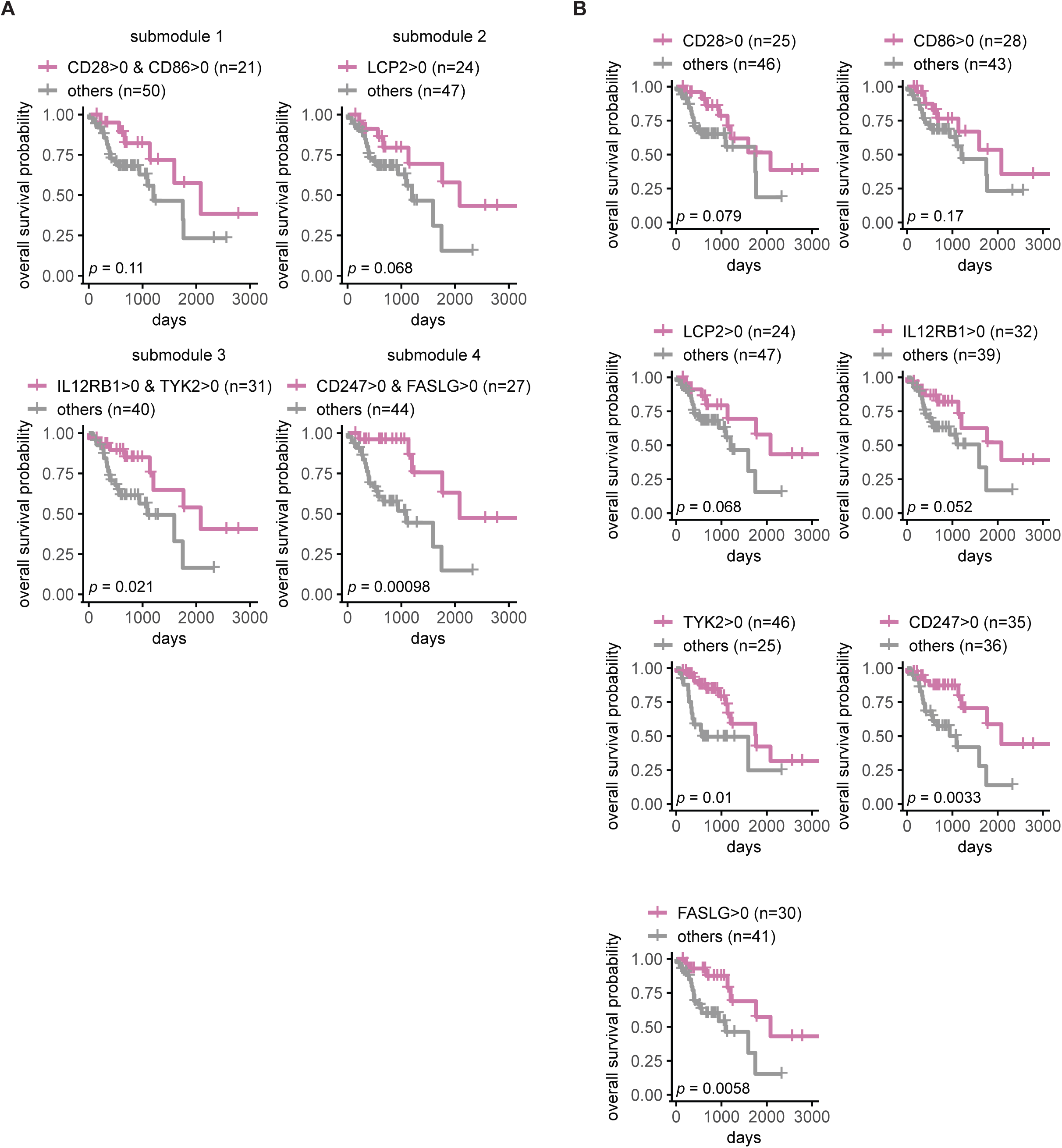
Overall survival of HPV+ exploratory set patient samples stratified by the inferred pathway levels (IPLs) of proteins from the same submodule (A) or individual protein (B). Not all overall survival tests were statistically significant under a log-rank *p* < 0.01 cutoff.

**Supplementary Figure 10.**
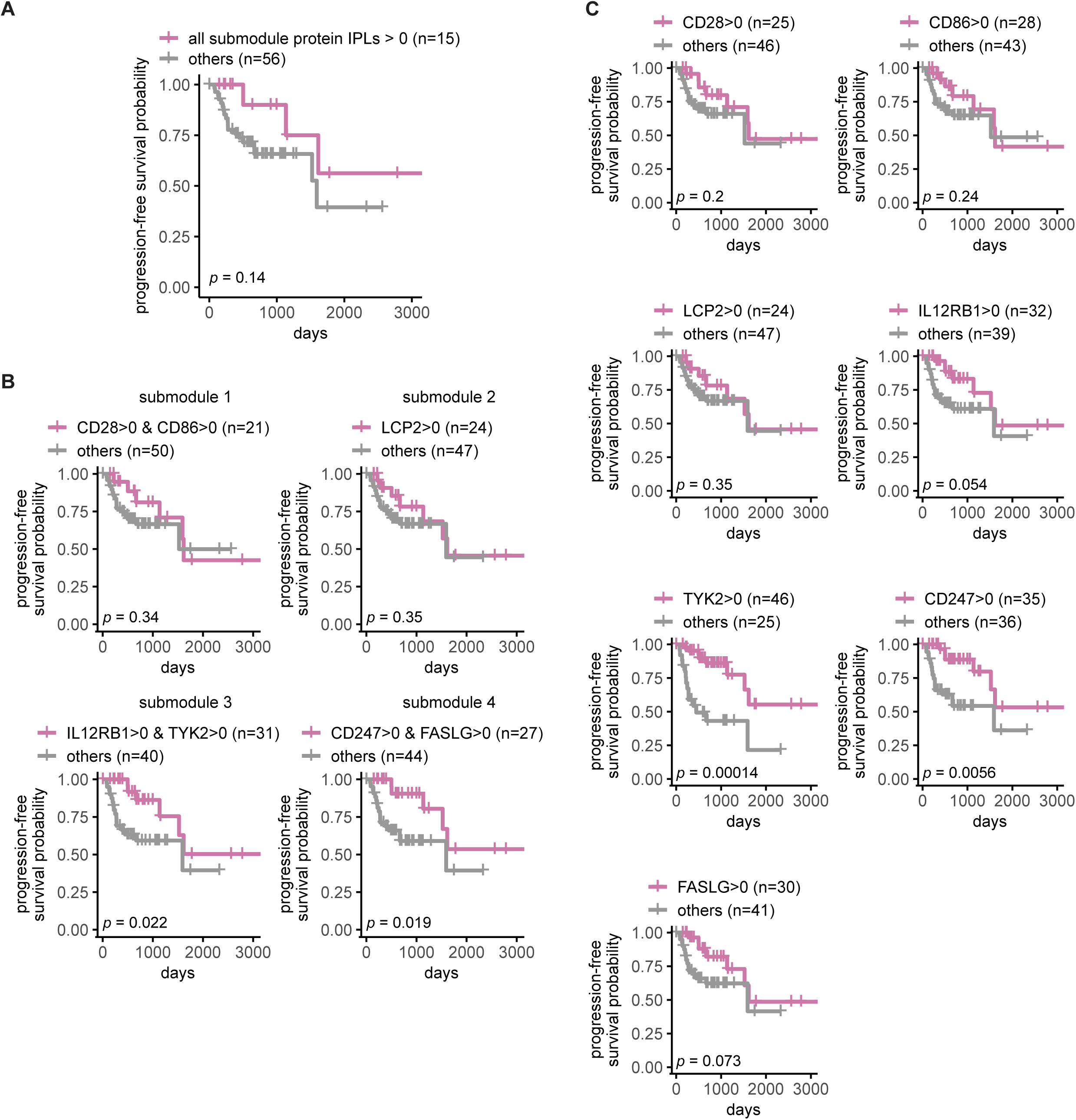
Progression-free survival of HPV+ exploratory set patient samples stratified by the inferred pathway levels (IPLs) of all the seven proteins combined (A); proteins from the same submodule (B), or individual protein (C). Not all progression-free survival tests were statistically significant under a log-rank *p* < 0.01 cutoff.

**Supplementary Figure 11.**
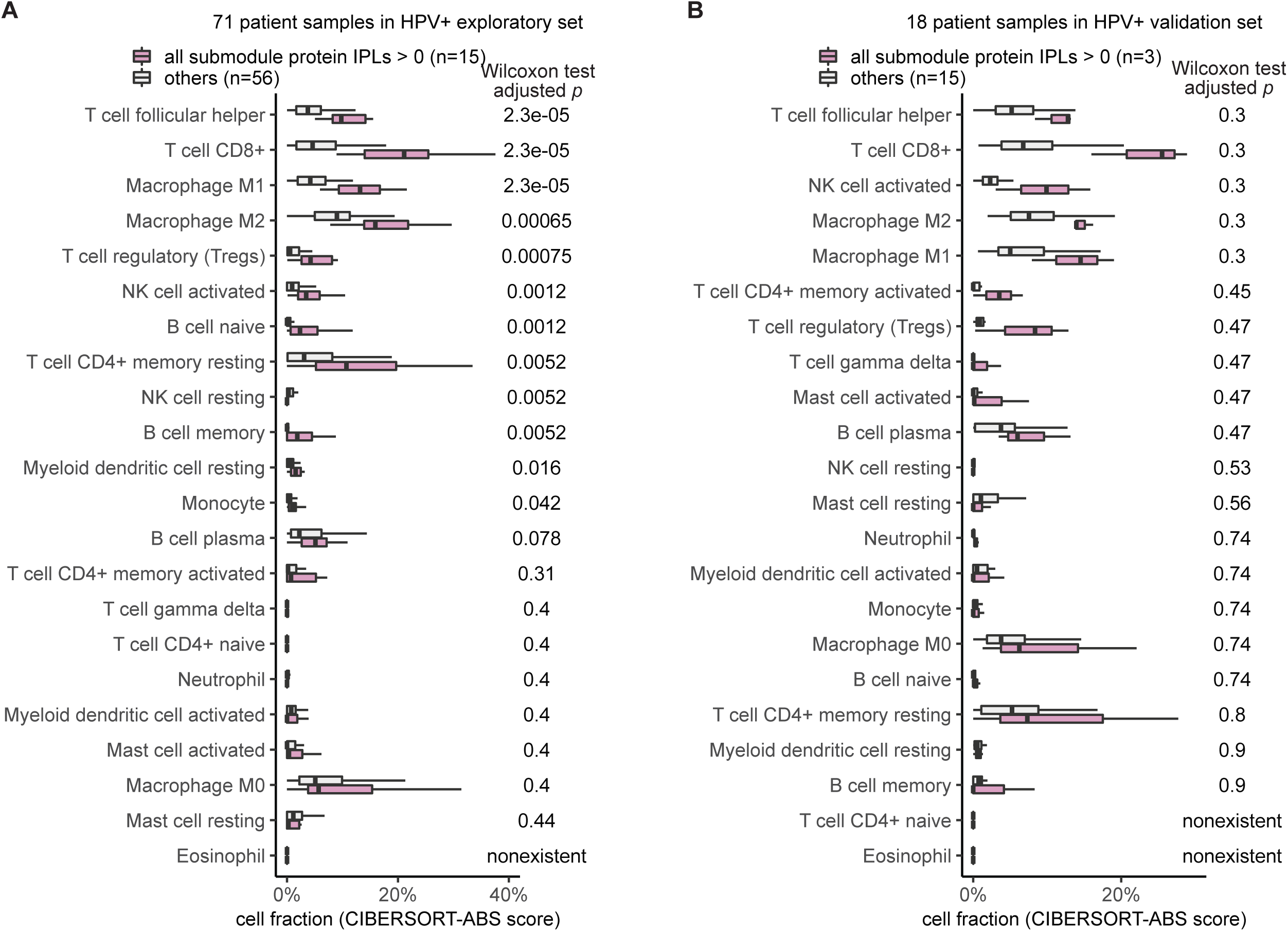
Immune cell compositions stratified by the inferred pathway levels (IPLs) of all the seven proteins combined for HPV+ exploratory (A) and validation (B) set. Adjusted Wilcoxon test *p* on the cell composition difference between two groups of patient samples were shown for each cell type. Eosinophil was not found in either the exploratory or validation sets. T cell CD4+ naïve was not found in the validation set.

**Supplementary Figure 12.**
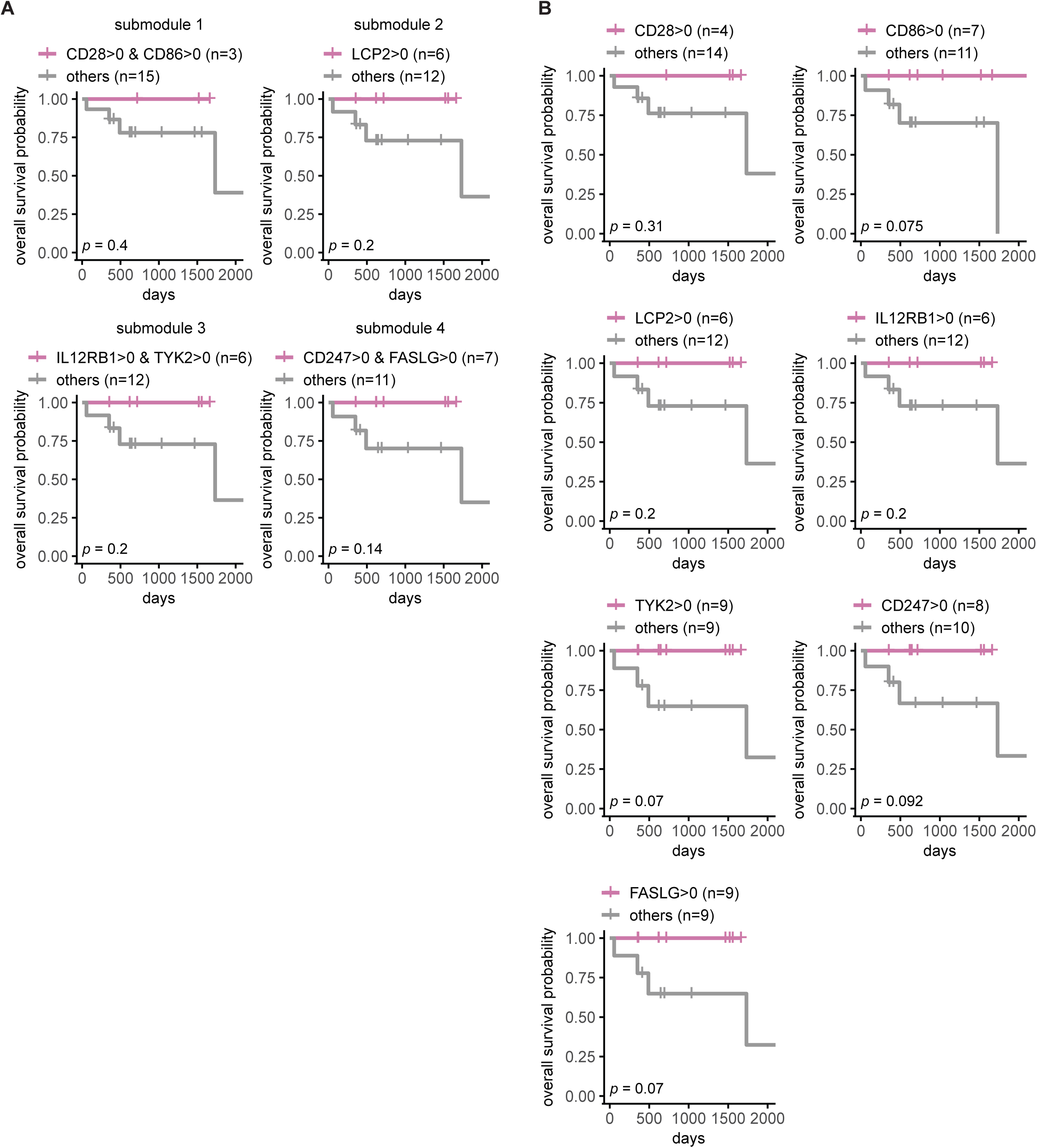
Overall survival of HPV+ validation set patient samples stratified by the inferred pathway levels (IPLs) of proteins from the same submodule (A) or individual protein (B). Not all overall survival tests were statistically significant under a log-rank *p* < 0.01 cutoff.

**Supplementary Figure 13.**
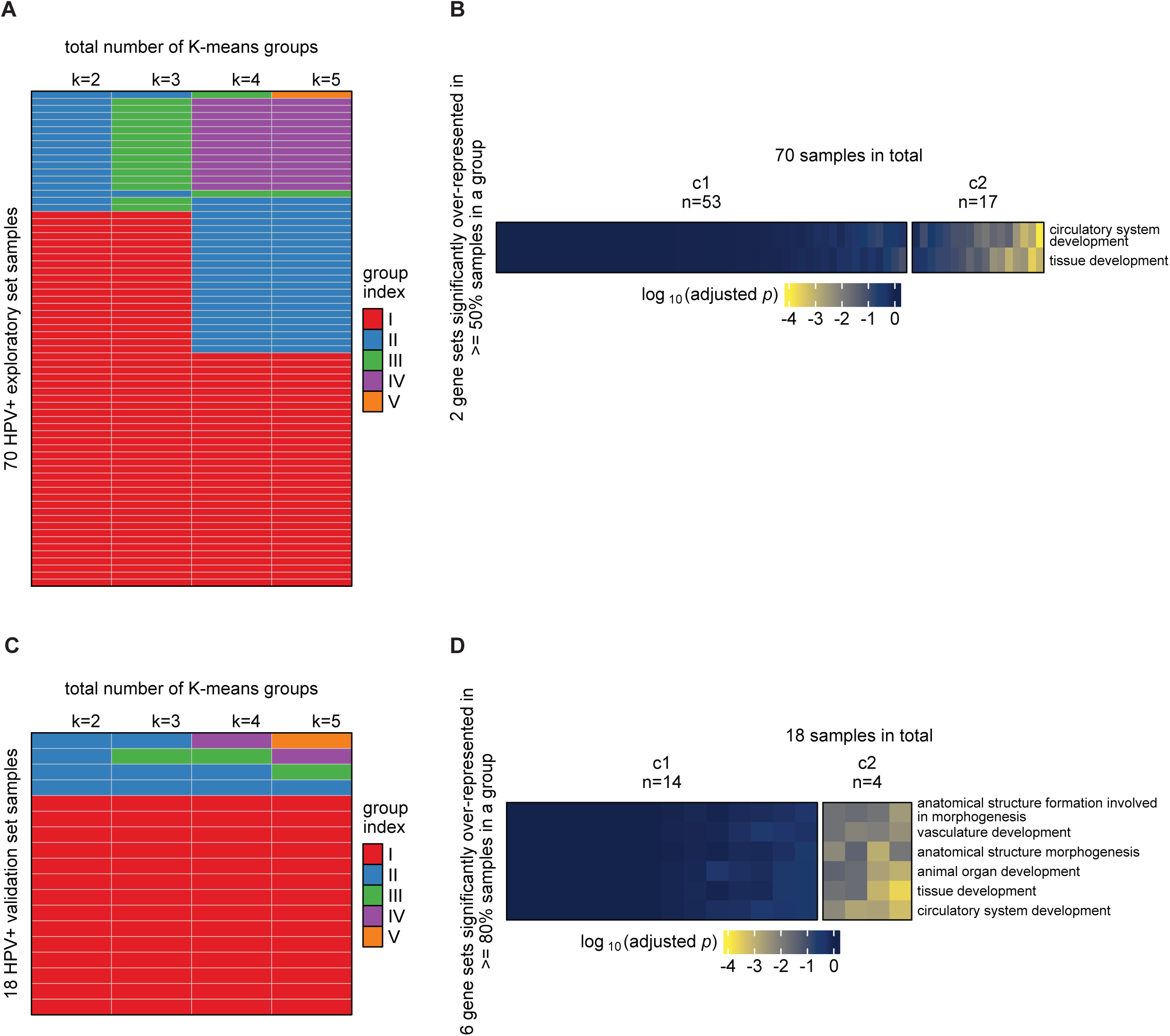
Clustering patient samples and annotating patient groups by PARADIGM IPLs from Pan-Cancer Atlas. K-means clustering applied to HPV+ exploratory (**A**) and validation (**C**) sets with a total number of groups ranging from two to five. Significantly overrepresented GO terms for groups of patients from HPV+ exploratory (**B**) and validation (**D**) sets. Note that one patient sample from HPV+ exploratory set does not have IPLs available in the Pan-Cancer Atlas’s PARADIGM results, so only 70 of the 71 samples are shown in **A** and **B**.

**Supplementary Figure 14.**
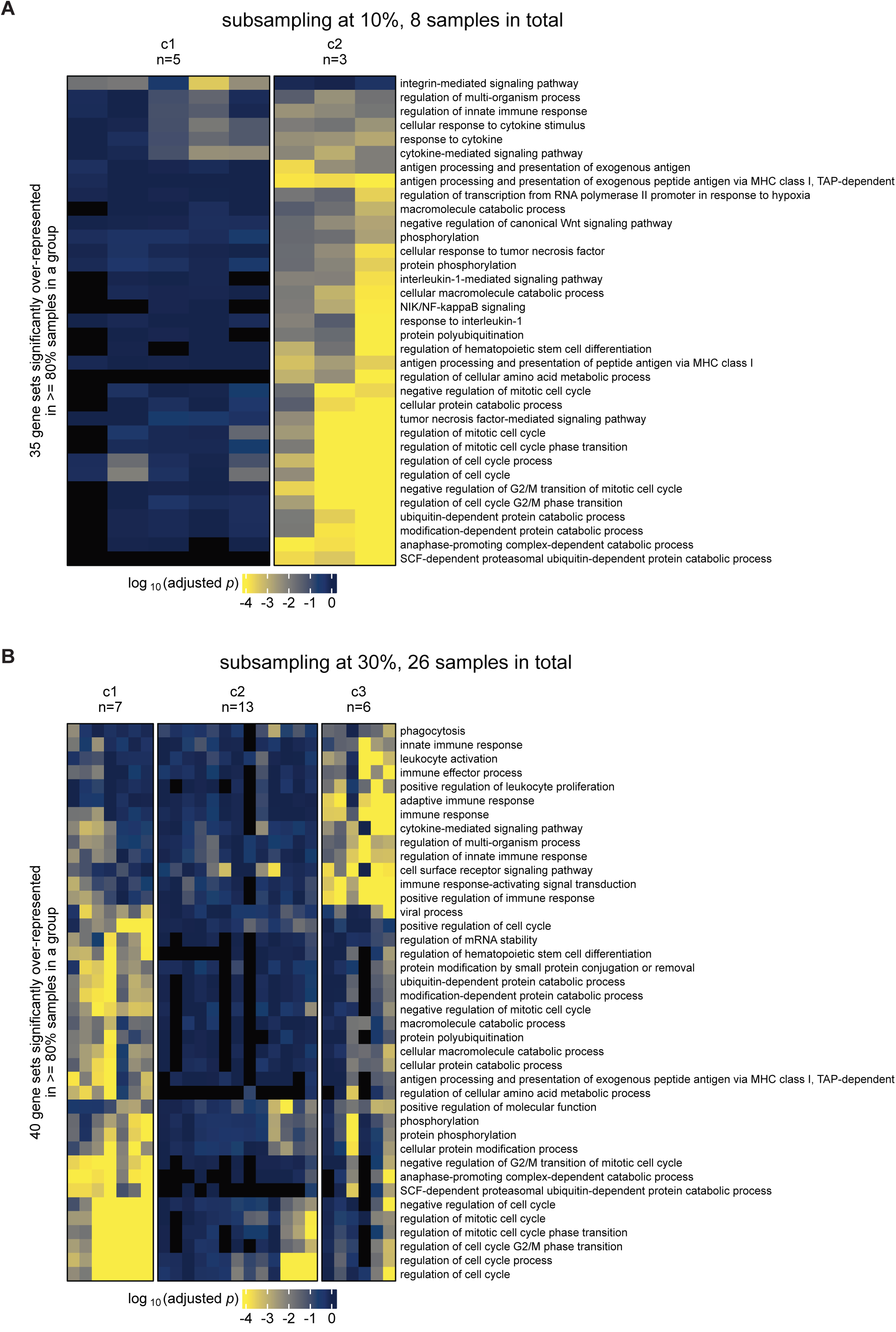
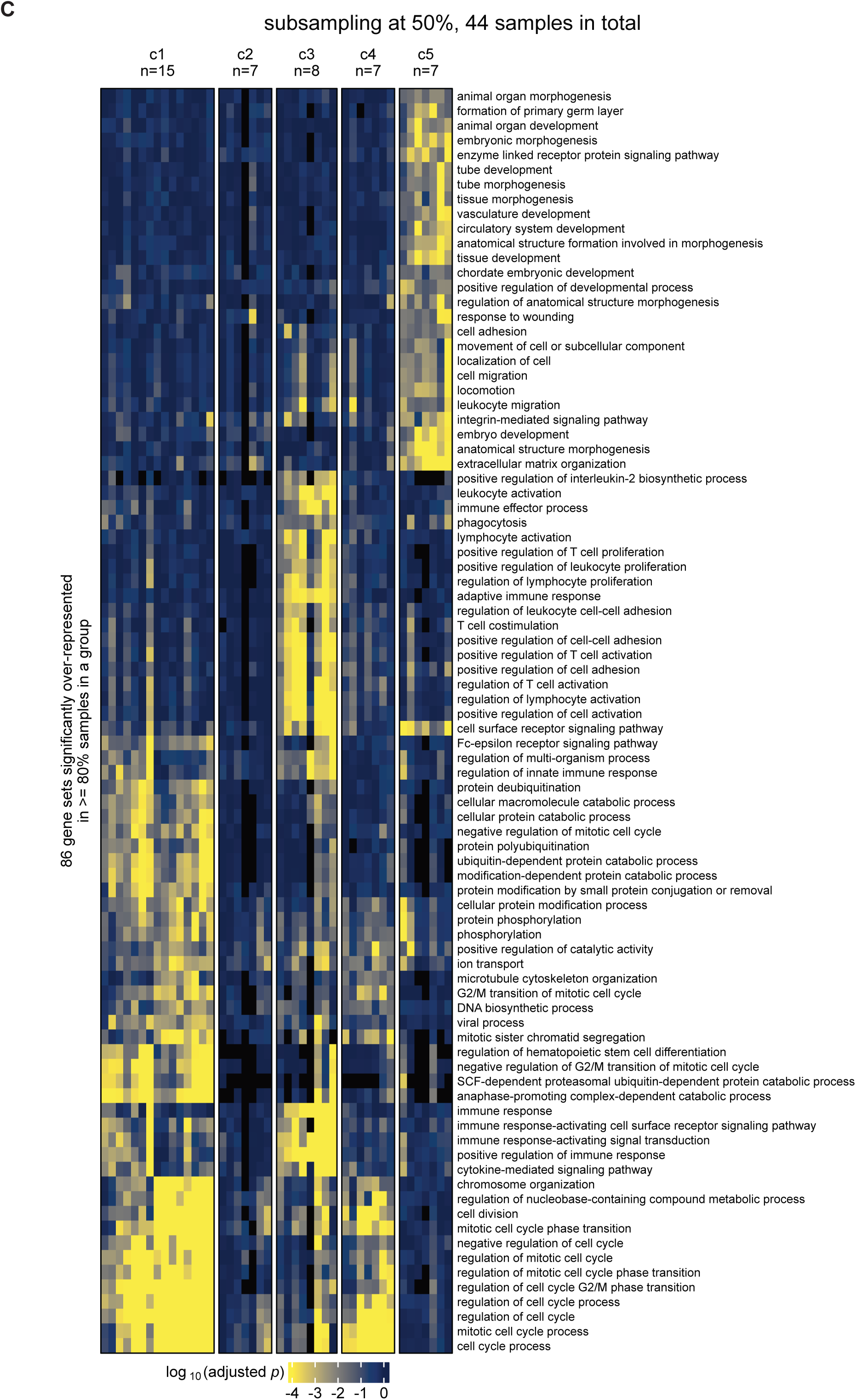
Sub-sampling analysis by MPAC on HPV+ patient samples. Patient groups and their significantly overrepresented GO terms by MPAC on randomly selected 10% (**A**), 30% (**B**), and 50% (**C**) of the 89 samples from HPV+ exploratory and validation set combined.

**Supplementary Figure 15.**
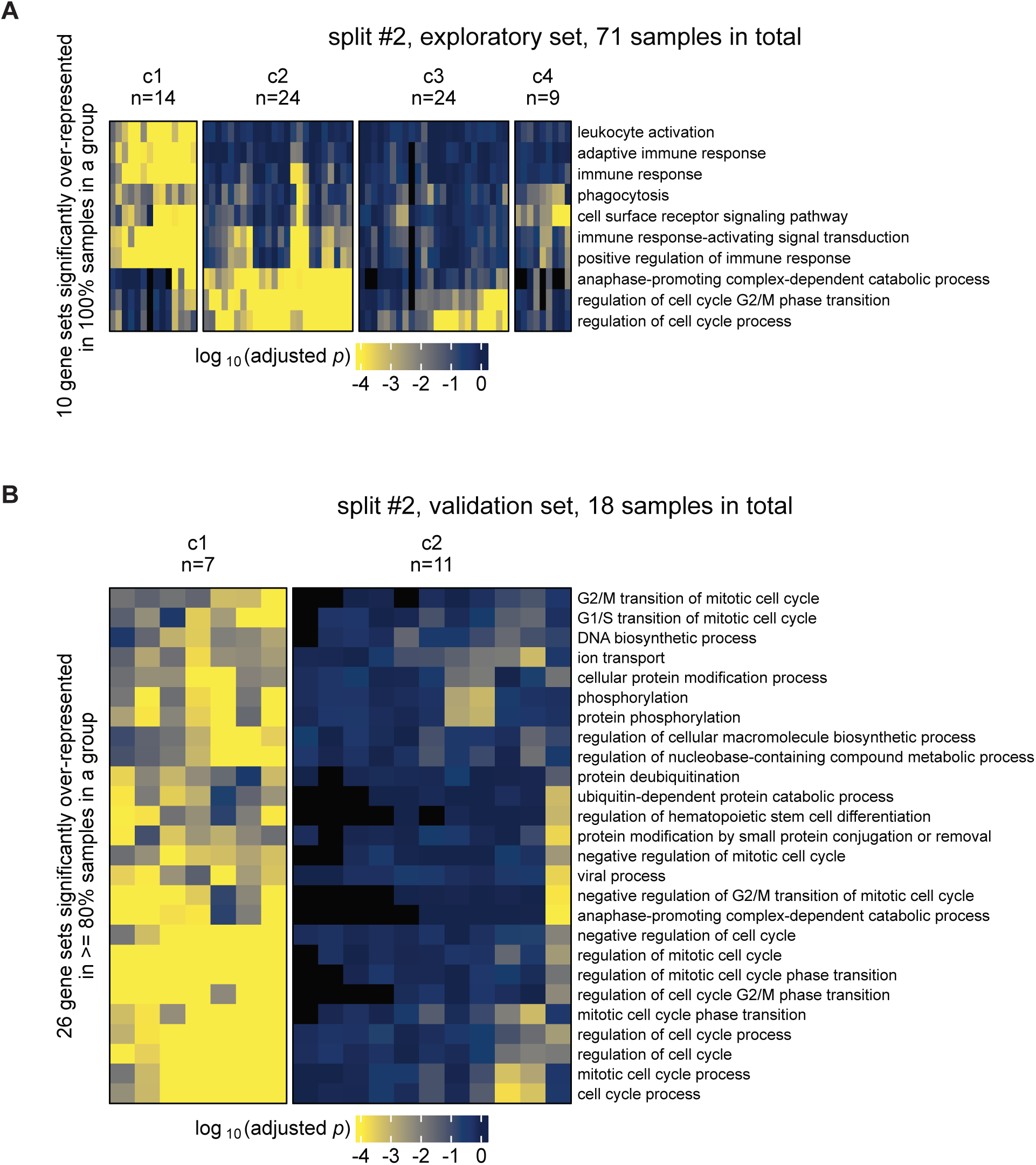
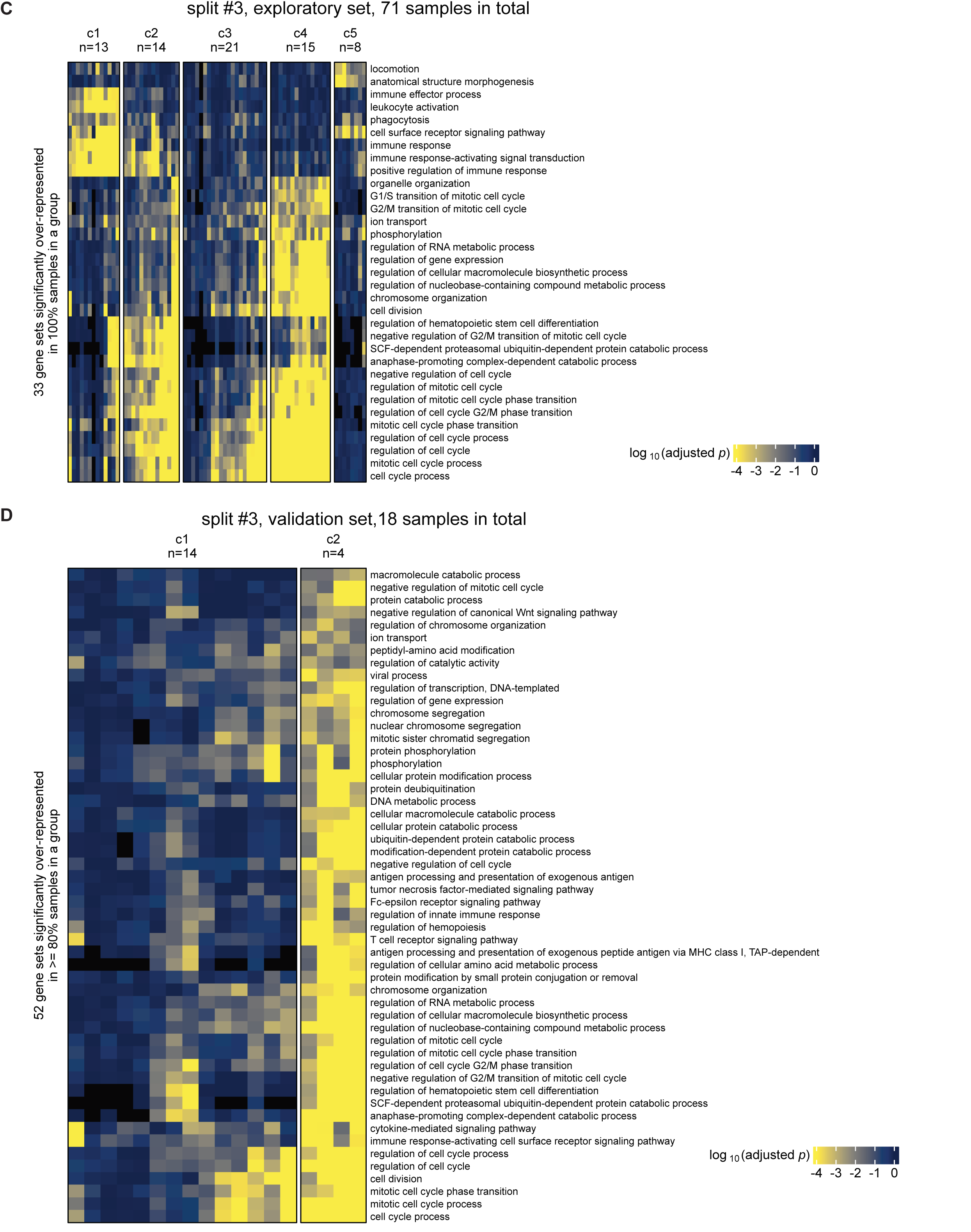
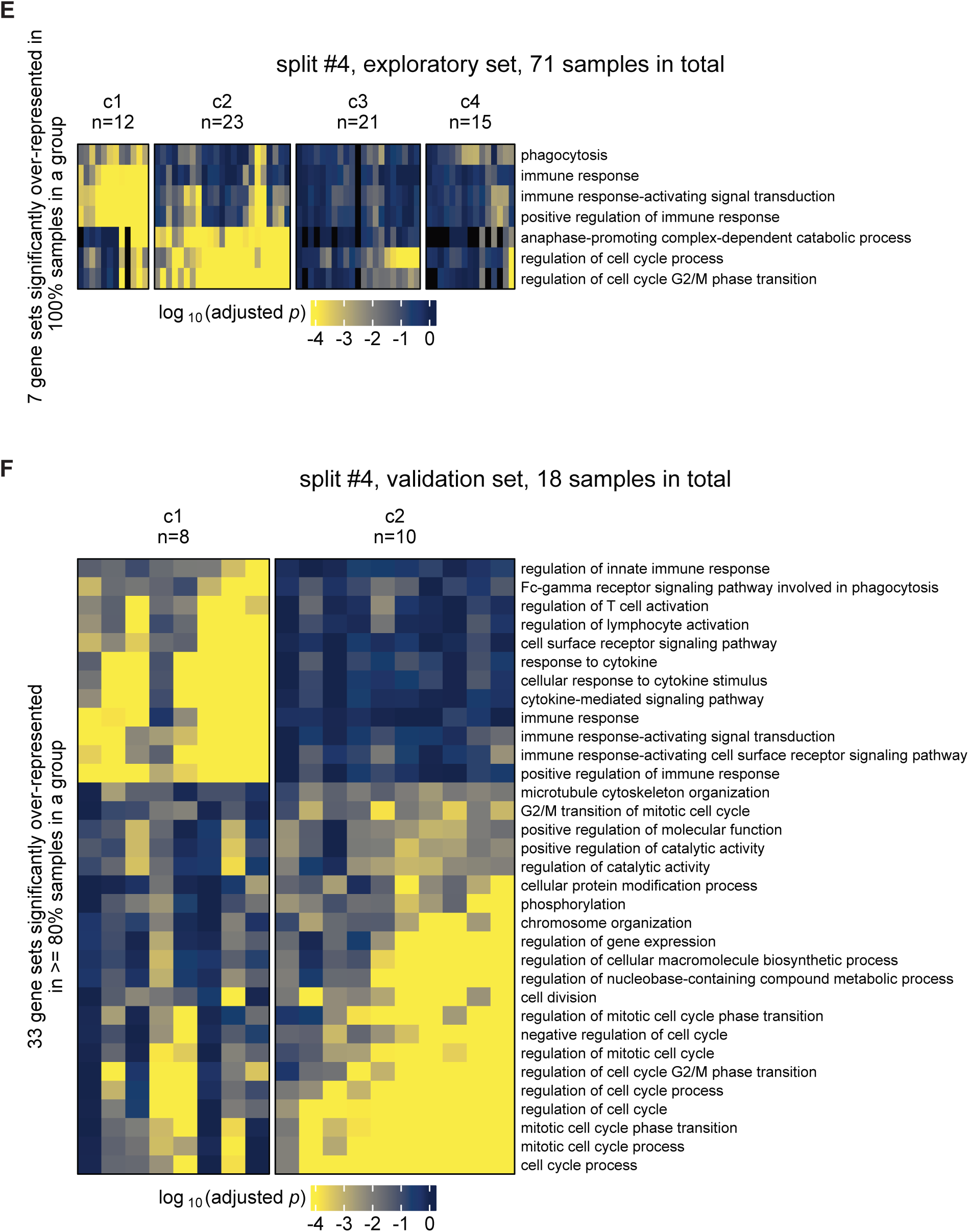
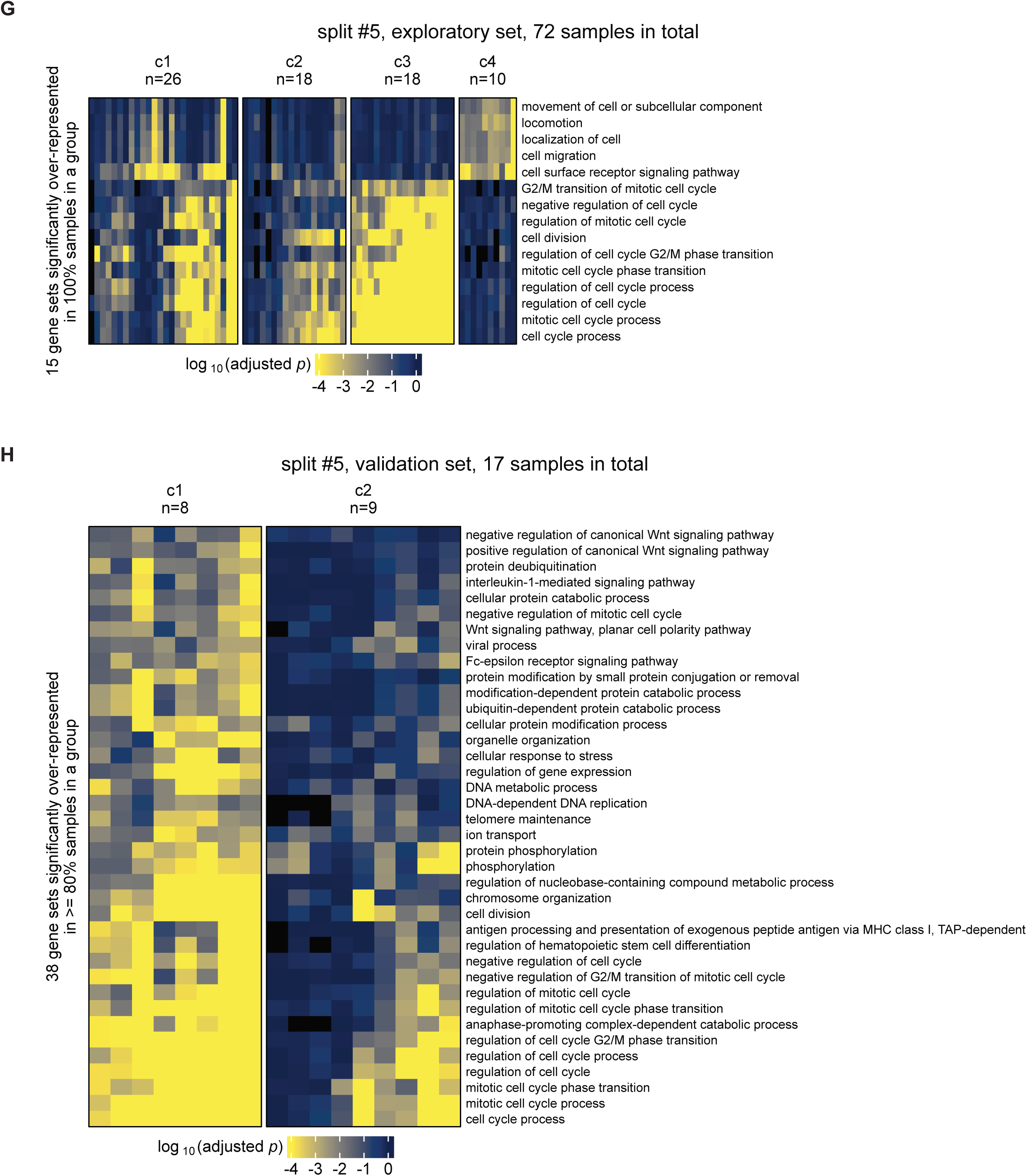
Exploratory and validation set resampling of MPAC on HPV+ patient samples. The 71 exploratory set samples were divided into four groups with similar sizes (18, 18, 18, and 17). The 18 validation set samples were used as the fifth group. The existing exploratory set and validation set was defined as split #1. Results for split #1 are shown in Figure 2 and Figure 4A. Patient groups and significantly overrepresented GO terms by MPAC are shown here for split #2 (**A** & **B**), #3 (**C** & **D**), #4 (**E** & **F**), and #5 (**G** & **H**).

**Supplementary Figure 16.**
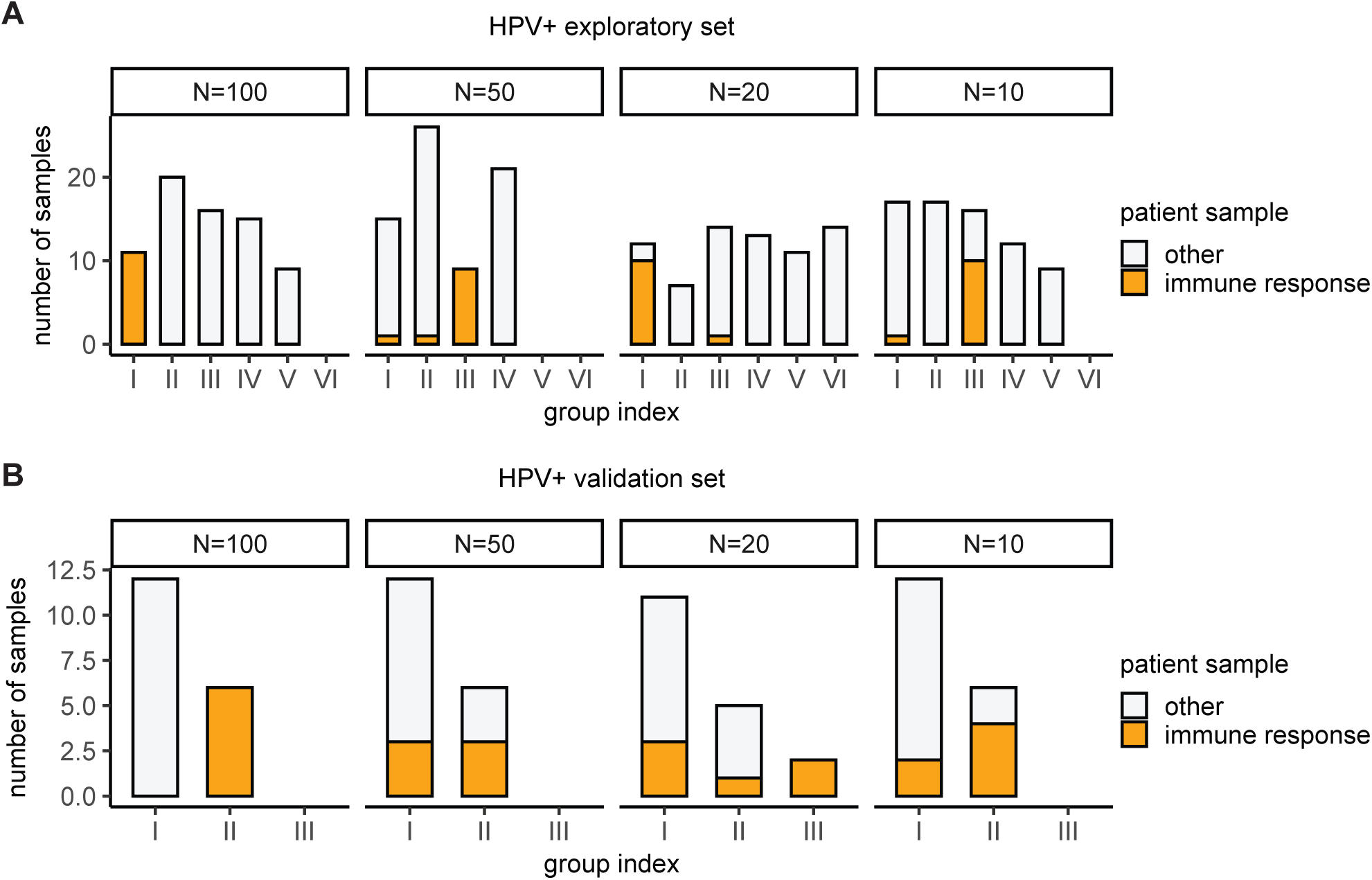
Distribution of immune response patient samples under different numbers of permutations for HPV+ exploratory (A) and validation (B) set. Immune response patient samples are defined as the eleven Group I exploratory set samples (Figure 2) and the six Group II validation set samples (Figure 4) from 100 permutations.

**Supplementary Figure 17.**
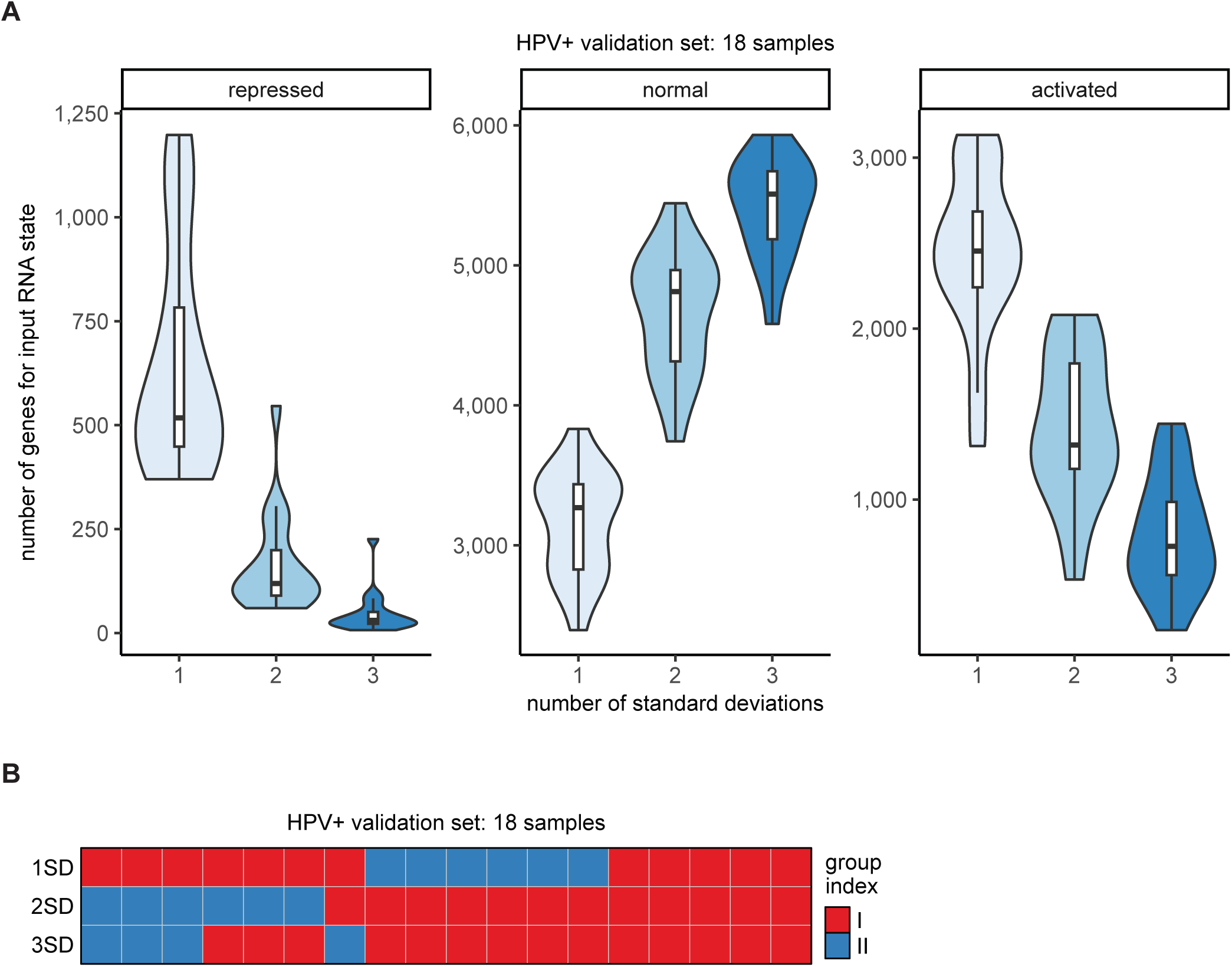
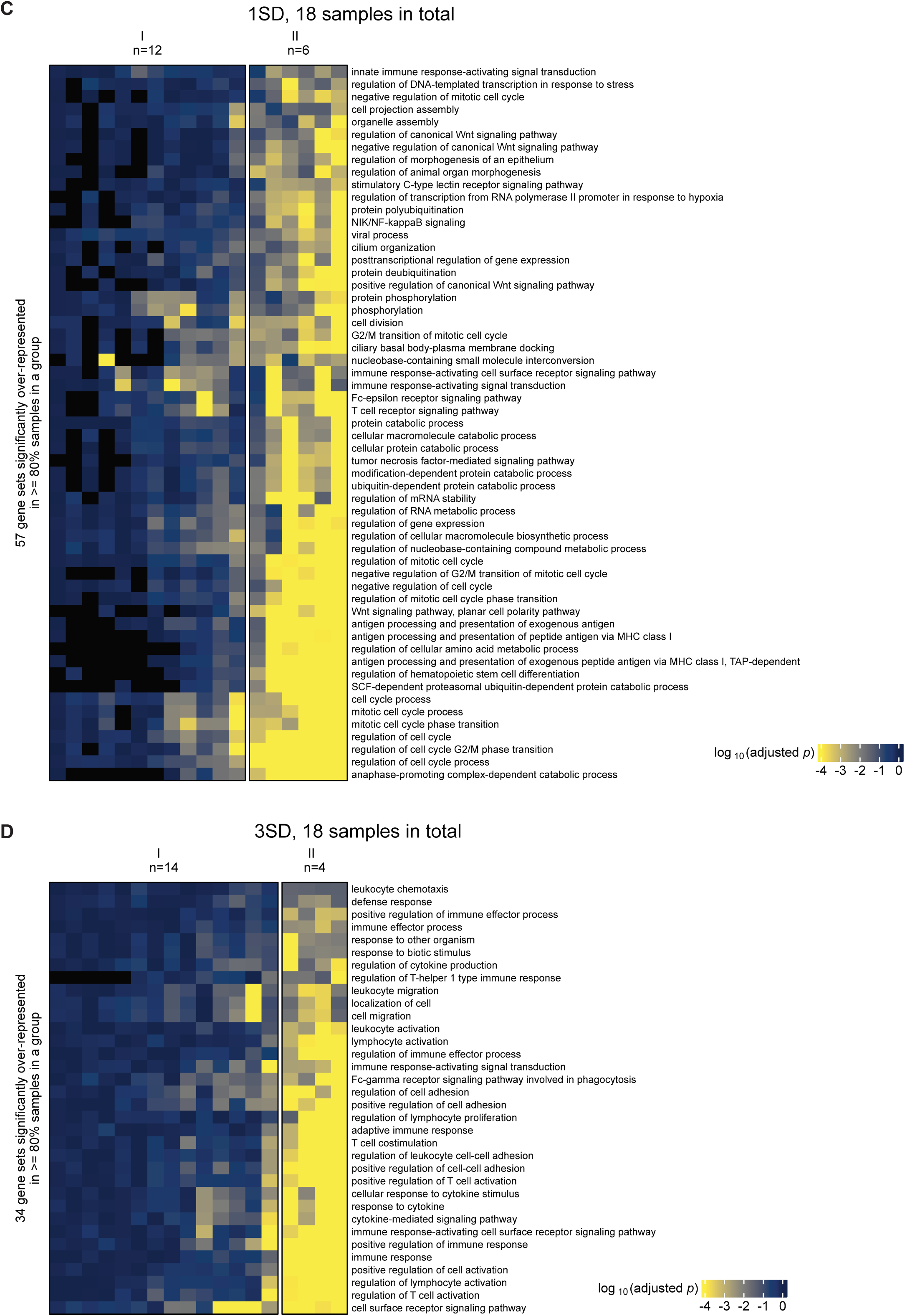
Impact of one, two, or three standard deviation thresholds on defining input RNA states for HPV+ validation set. (**A**) Distributions of input RNA states; (**B**) Patient groups; (**C** & **D**) Significantly overrepresented GO terms in each patient group under one (**C**) or three (**D**) standard deviation threshold. Those under two standard deviations are shown in Figure 4A.

**Supplementary Figure 18.**
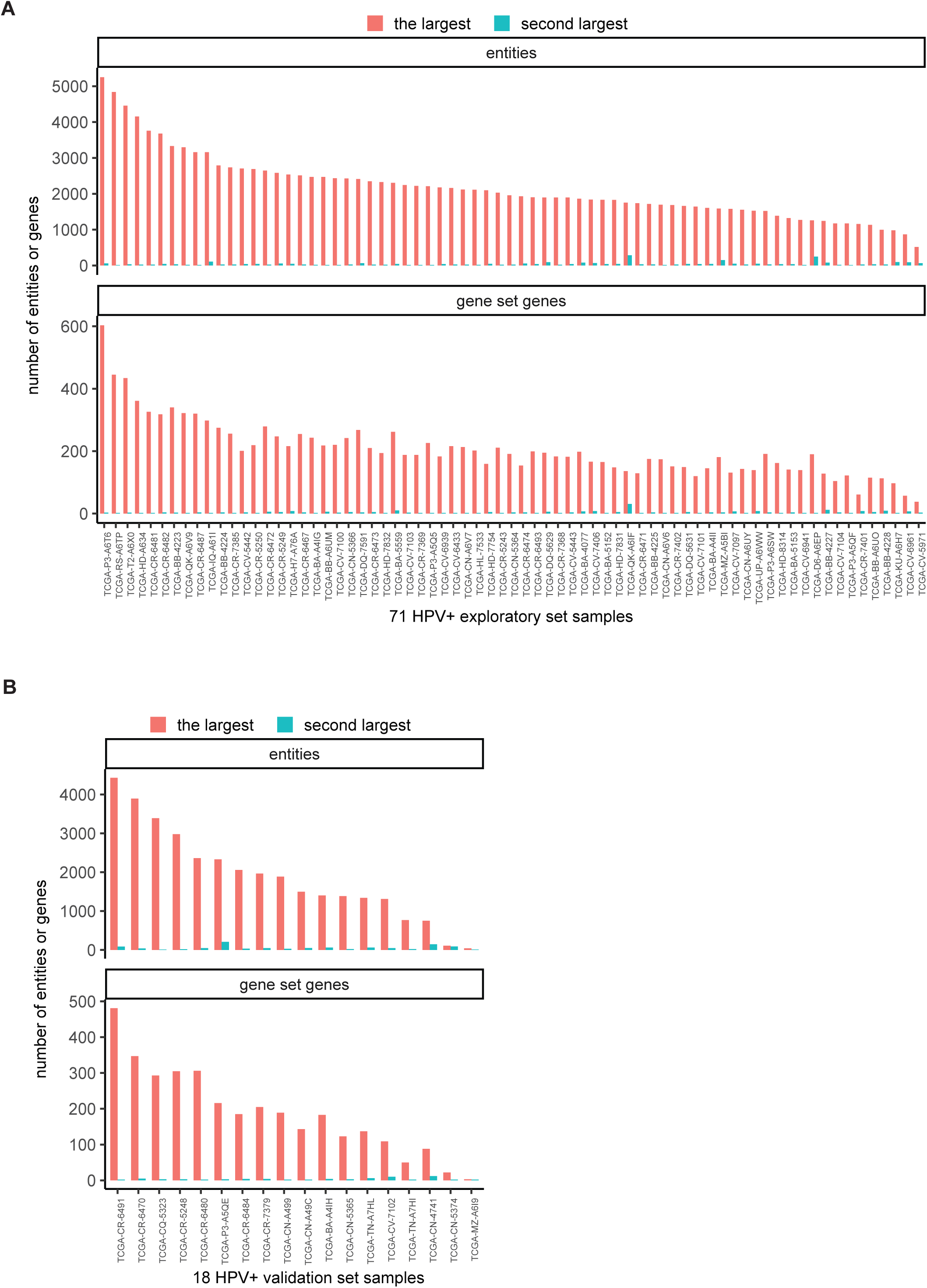
Comparison of the sizes of the largest and the second largest sub-network for HPV+ exploratory (A) and validation (B) set. Sizes are represented by the total number of entities or the number of gene set genes that were used for downstream GO term overrepresentation analysis.

**Supplementary Figure 19.**
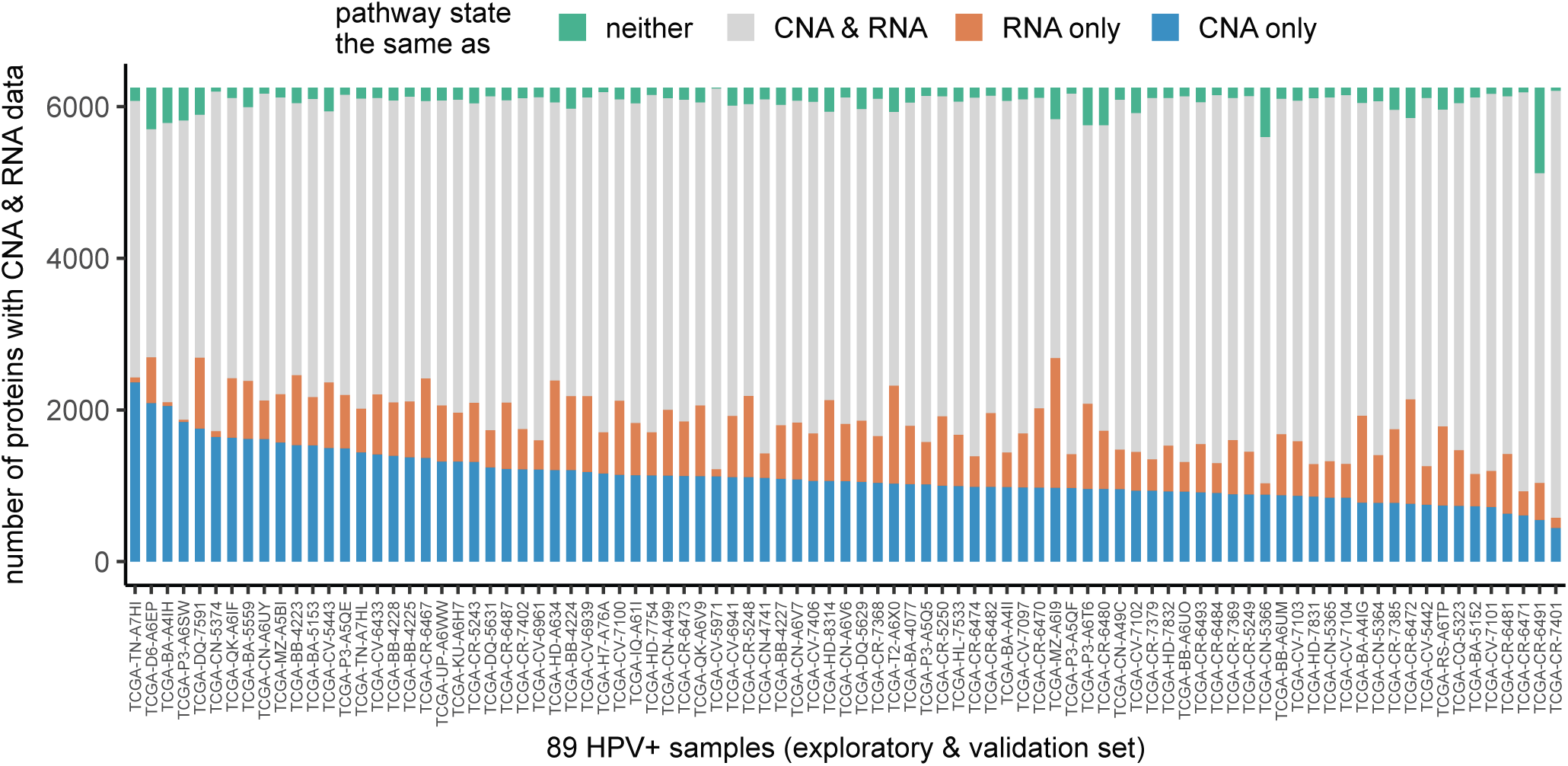
Comparison of the number of proteins that have their pathway states agreed with CNA and/or RNA-seq data. All the 6,251 proteins that have CNA and RNA-seq data were stratified by whether their pathway states agree with neither of their CNA or RNA states (green), both CNA and RNA states (gray), only RNA but not CNA states (orange), or only CNA but not RNA states (blue).

**Supplementary Figure 20.**
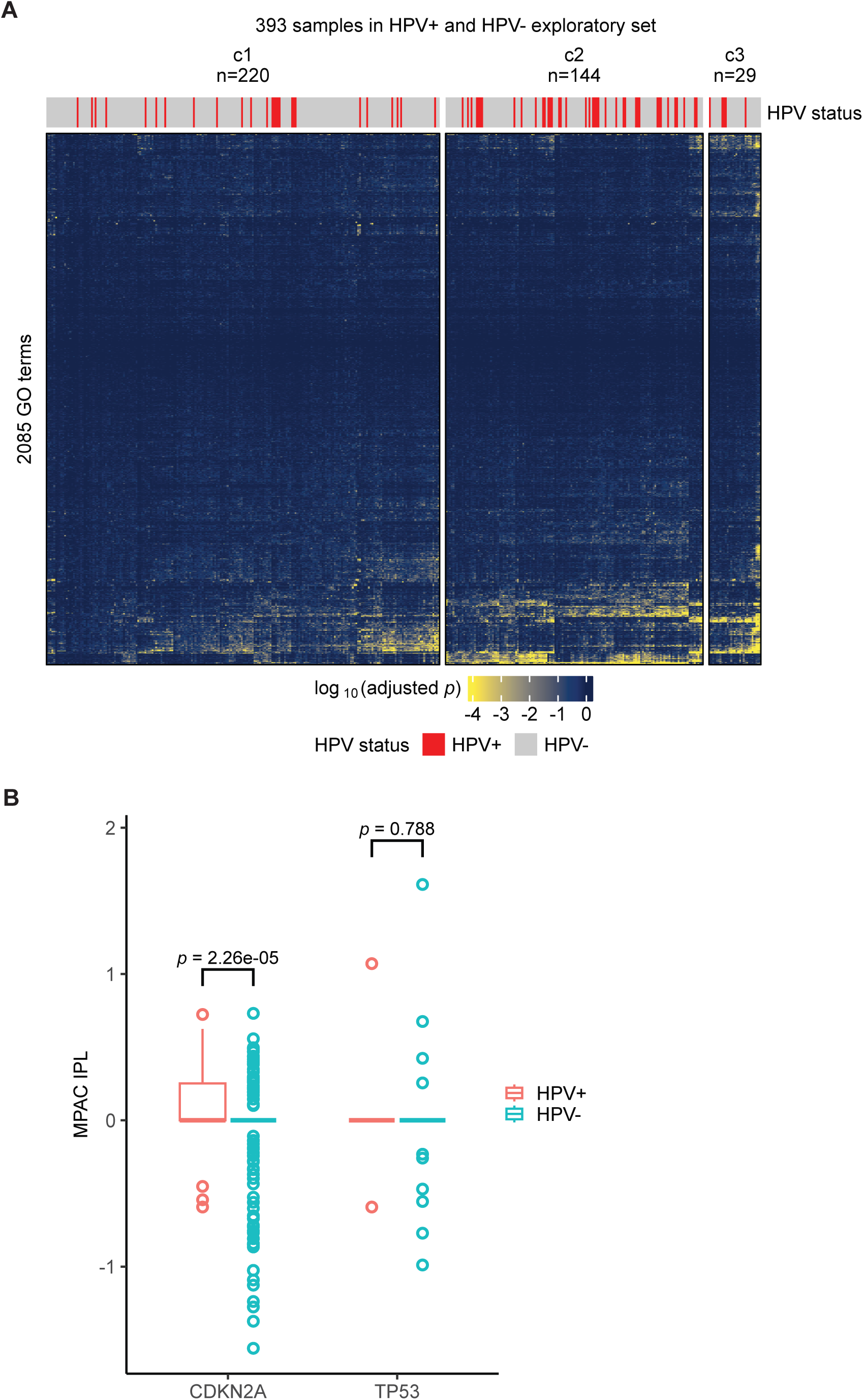
MPAC cannot separate HPV+ and HPV- samples due to insufficient input pathway knowledge. (**A**) MPAC grouping of patient samples from HPV+ (red) and HPV- (gray) exploratory set by their GO term overrepresentations; (**B**) Comparison of MPAC IPLs of two HPV subtype-specific proteins, CDKN2A and TP53, in patient samples from HPV+ (red) and HPV- (green) exploratory set. Wilcoxon *p*-values on the comparison are denoted.

**Supplementary Figure 21.**
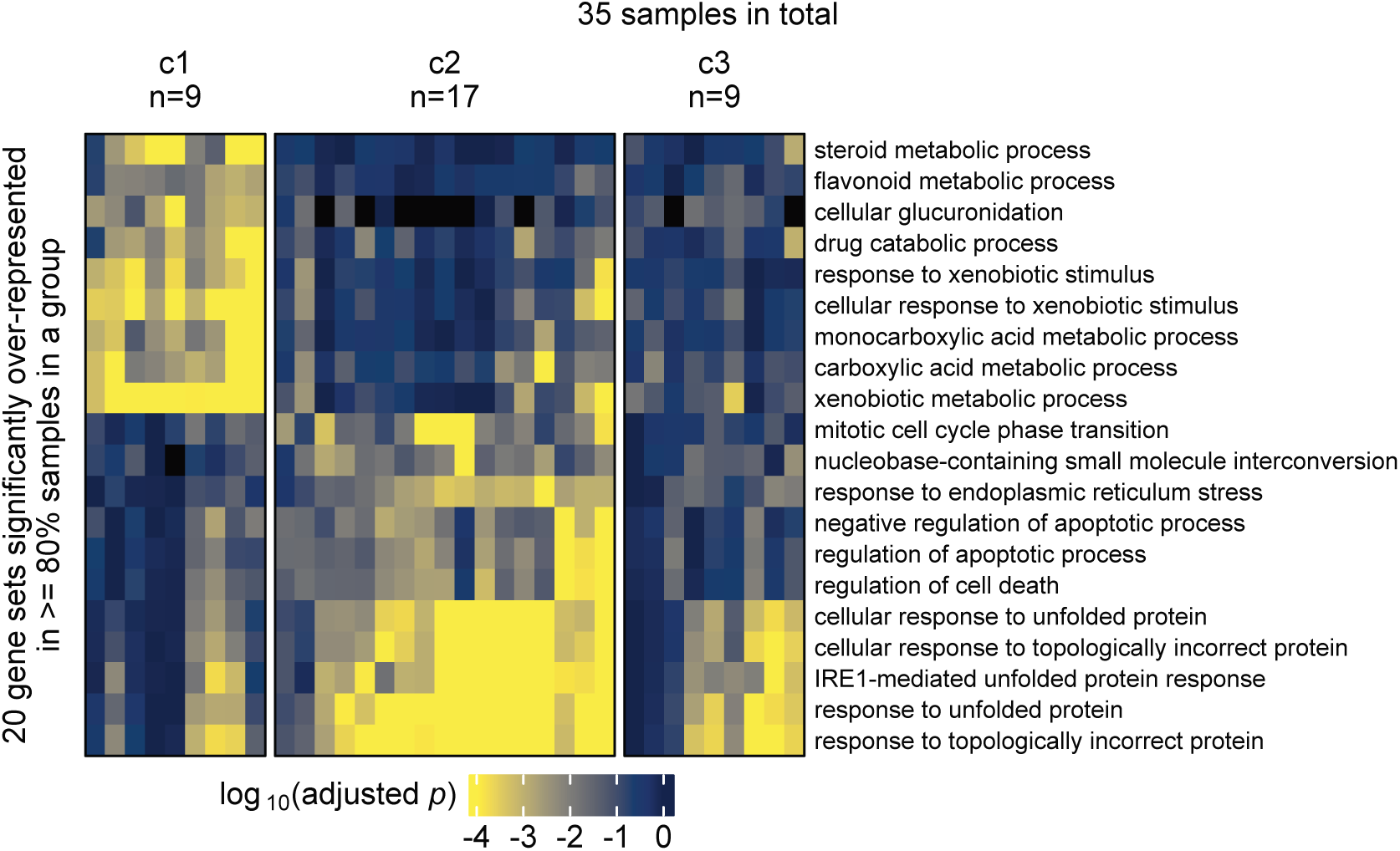
Patient sample groups and significantly overrepresented GO terms for the TCGA cholangiocarcinoma cohort.

**Supplementary Figure 22.**
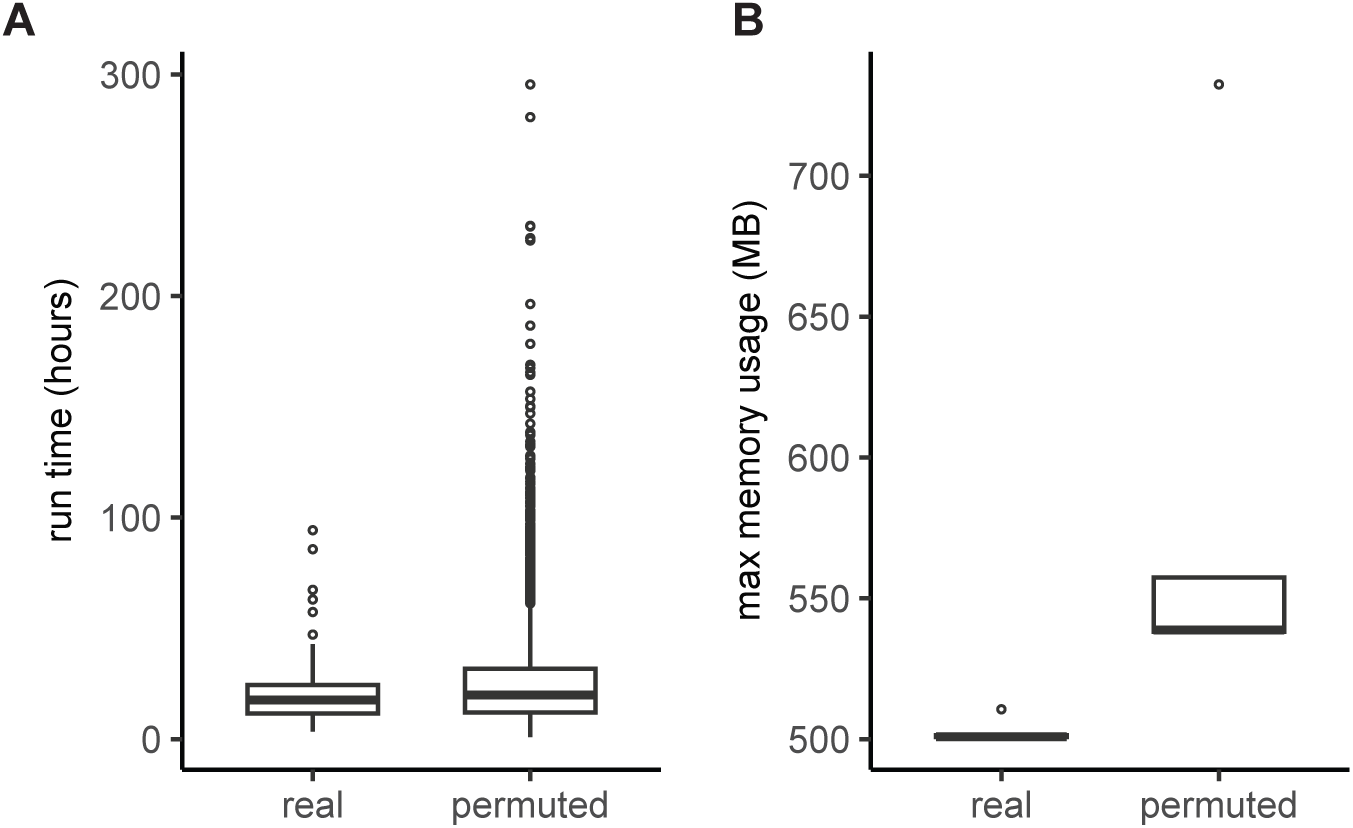
Time (A) and memory (B) usage by MPAC’s PARADIGM subroutine runs on real and permuted data from the 71 HPV+ exploratory set patient samples. All the jobs were run via HTCondor on machines from the UW–Madison Center for High Throughput Computing.

### Supplementary Notes

**Supplementary Note 1. Pseudocode of MPAC.**

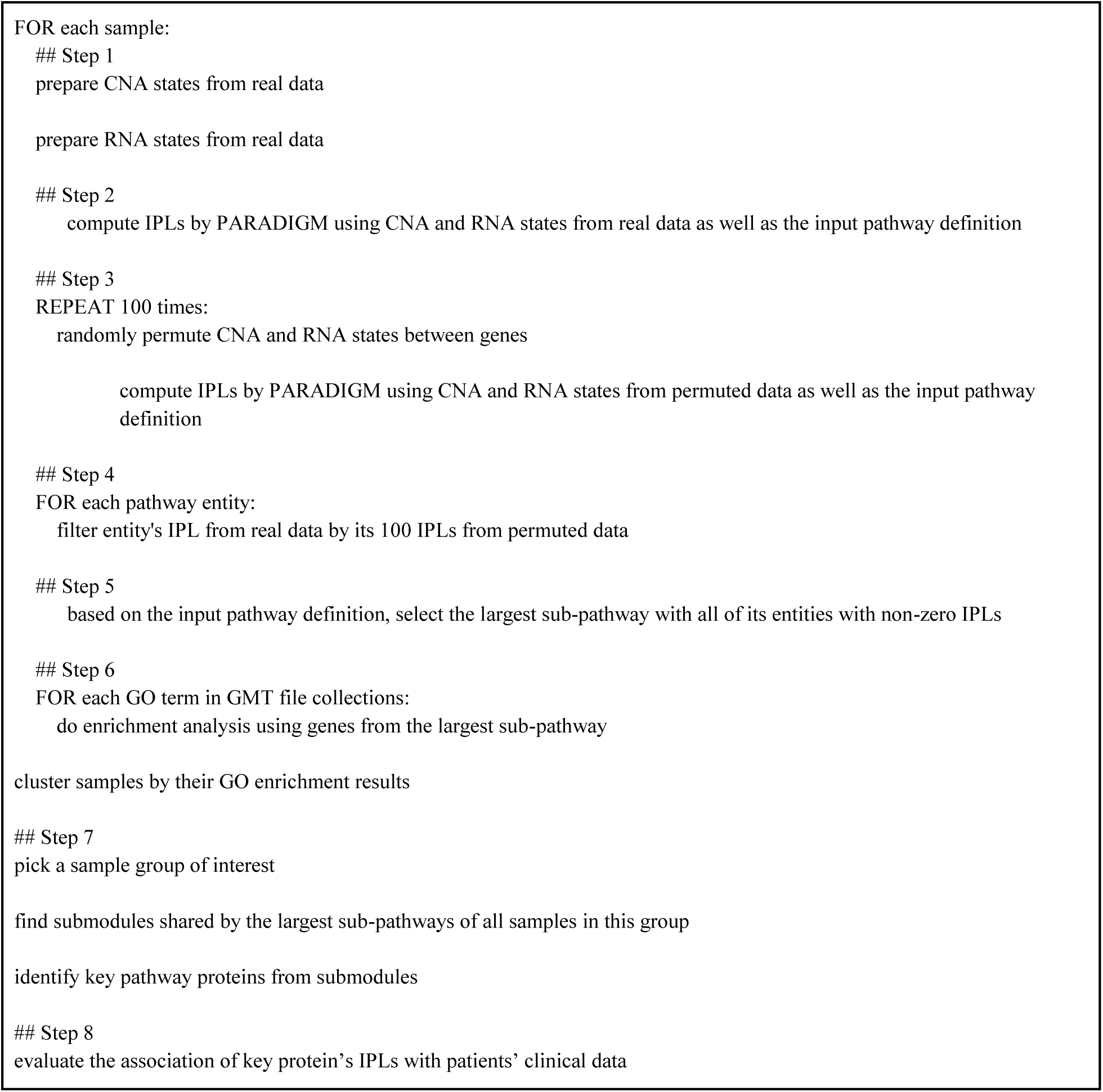

**Supplementary Note 2. Required number of permutations in MPAC.** To find out the number of permutations required for a valid result, we randomly selected 10, 20, and 50 permutations from the existing 100 permutations for each HPV+ sample. With them, we ran MPAC under the same protocol. We checked if the 11 and 6 immune response samples from the exploratory and validation set, respectively, were still grouped together and with no other samples added under a smaller number of permutations. For the exploratory set, when using 50 or 20 permutations, 9 or 10 of the 11 immune response samples continued to fall into the same group with at most 1 other sample added (Supplementary Figure 16A). When using 10 permutations, although 10 immune response samples were clustered together, 6 other samples came to the same group (Supplementary Figure 16A) and thus added more noise to the biological function of this group. For the validation set, the 6 immune response samples were spread almost evenly into different groups with at most 3 or 4 samples in a group with 2 or more other samples added by 50, 20, or 10 permutations (Supplementary Figure 16B). This analysis suggests that if a user has about 70 samples like the HPV+ exploratory set, 50 or 20 permutations would be ok. If a user has about 20 samples like the HPV+ validation set, 100 permutations would be required. In the MPAC package, we set the default to 100 and provide an option for users to adjust this number based on their sample sizes.

**Supplementary Note 3. Number of standard deviation threshold to define input RNA states.** The reason MPAC takes two standard deviations (SD) as the threshold is because it corresponds to the commonly used *p* < 0.05 cutoff. For any gene, MPAC extracts its expression levels from RNA-seq data of normal samples and fits them by a Gaussian distribution. Two-SD threshold on such a Gaussian distribution is at a *p* = 0.05 cutoff (three SD is at a *p* = 0.01 cutoff and one SD is at a *p* = 0.32 cutoff). To evaluate the impact by different SD, we tried 1, 2, and 3 SD on the 18 samples in the HPV+ validation set. We first checked their effects on the RNA states of all the genes. Those from 2 and 3 SD are similar to each other, especially for repressed and normal states (Supplementary Figure 17A), while those from 1 SD are largely different from those of 2 or 3 SD. Next, we compared the sample groups from different SD. Groups from 2 and 3 SD share good overlaps with three Group II samples in common (Supplementary Figure 17B), whereas groups from 1 SD are totally different from those from 2 and 3 SD. Such dissimilarity is further supported by the overrepresented pathways for each group. By 1 SD, its Group II has many GO terms related to cell cycle, Wnt signaling, etc, but very few related to immune response (Supplementary Figure 17C). In contrast, by 3 SD, its Group II has a large fraction of overrepresented GO terms related to immune response (Supplementary Figure 17D), which is similar to those by 2 SD (Figure 4A). Taken together, because of the commonly used *p* < 0.05 or < 0.01 cutoff, and because of the agreement on the RNA states and sample groups, we recommend users consider 2 or 3 SD to define input RNA states.

**Supplementary Note 4. The largest pathway sub-networks are much bigger than the second largest ones.** In MPAC, each sample has its own largest sub-network. The reason why MPAC only considers the largest sub-network is because the second largest sub-network is much smaller compared to the largest one. To illustrate this, we compared the sizes between the two from each sample in terms of two types of entities: (1) all entities in the sub-networks, which include proteins, complexes, families, etc; (2) only proteins that have their genes in the GO terms that were used for MPAC’s enrichment analysis. In both HPV+ exploratory and validation set, the largest sub-networks are overwhelmingly larger than the second largest sub-networks for both all entities and GO terms genes (Supplementary Figure 18). The sub-network’s ‘stability’ could be affected by a few factors, such as the number of permutations or the number of standard deviations to define input RNA states.

**Supplementary Note 5. CNA has a stronger impact than RNA-seq in determining protein pathway states.** MPAC integrates both CNA and RNA-seq data, but CNA’s role is not evident from the key proteins shown earlier (Supplementary Figure 5-7). We extended the analysis to all the 6,251 pathway proteins with both CNA and RNA-seq data for all the 89 HPV+ exploratory and validation set samples. The comparison results are divided into the following four categories by whether a pathway state:

- Different from the one of CNA and RNA
- Same as the one of both CNA and RNA
- Same as the one of RNA, but different from the one of CNA
- Same as the one of CNA, but different from the one of RNA

For the majority of the 89 samples, the fraction of proteins that have pathway states only the same as CNA (blue bars in Supplementary Figure 19) is higher than those that have pathway states only the same as RNA (orange bars in Supplementary Figure 19), indicating that CNA has a stronger impact than RNA on determining a protein’s pathway state overall. We also note that CNA does not have a stronger impact than RNA in every sample. 10 of the 89 samples have RNA with a stronger impact than CNA.

**Supplementary Note 6. MPAC cannot separate HPV+ and HPV- patients with insufficient input pathway information.** The reason we split HPV+ and HPV- before applying MPAC is because they are largely different in oncogenic pathways, tumor biology, and clinical treatment response. To test if MPAC can separate HPV+ and HPV-, we applied MPAC on the 393 samples from the HPV+ and HPV- exploratory sets. MPAC gives three groups and every group contains both HPV+ and HPV- samples (Supplementary Figure 20A), indicating that MPAC cannot separate HPV+ and HPV-. This is largely because the input pathways for MPAC have insufficient information on HPV. The pathways are from TCGA Pan-Cancer Atlas and are not designed specifically for HPV. They do not have any information on HPV protein E7 and very little on HPV protein E6. For example, it is well known that HPV E6 represses TP53 in HPV+, but MPAC’s pathways do not contain such interaction. To illustrate further, we checked the MPAC IPLs of TP53 as well as another protein CDKN2A, which is known to have alterations predominantly only in HPV-. For both of them, their IPLs do not show a difference between HPV+ and HPV- (Supplementary Figure 20B). Despite there being a statistical difference of CDKN2A IPLs (Wilcoxon *p*=2.26×10^-5^), CDKN2A and TP53’s IPLs are within the same IPL ranges between HPV+ and HPV-, illustrating the impact of insufficient input pathway knowledge on key HPV-specific proteins and the difficulty on separating HPV+ and HPV-. The TCGA HNSCC study does not separate HPV+ and HPV- either using PARADIGM alone. In their paper’s Supplementary Figure S7.11 (https://doi.org/10.1038/nature14129), HPV+ samples are mainly in subtype 3 (black), but this subtype also contains a substantial number of HPV- samples.

**Supplementary Note 7. Time and memory requirements for MPAC.** The major computational bottleneck for MPAC is running PARADIGM using the Pan-Cancer Atlas pathways on the large number of permuted data. For the 71 samples in HPV+ exploratory set, 100 permuted data for each sample resulted in 7,100 PARADIGM runs. This is the main reason why we utilized HTCondor from the Center for High Throughput Computing. Although most jobs on permuted data finished in 3 days, the longest one took about two weeks (Supplementary Figure 22A). In comparison, for jobs on real data, all of them finished in about four days (Supplementary Figure 22A). These jobs’ memory usage is not heavy. Only 500 to 600 MB is needed for most of them for both real and permuted data (Supplementary Figure 22B). Jobs on permuted data may require longer time and more memory than on real data because permuted data likely have input omic states that are discordant with the input pathways and thus it is more computationally expensive to find optimal pathway states for both. Lastly, please note that, because of the nature of HTCondor, the aforementioned jobs were run on a heterogeneous resource (e.g., CPU speed, architecture).

**Supplementary Note 8. Changes in newer versions of the igraph R package affected the reproducibility of our results but not the main conclusions from our analyses.** Starting in igraph version 1.3, the Louvain method is no longer deterministic and different runs may generate different clustering results (https://github.com/igraph/rigraph/issues/539). To evaluate the impact of non-deterministic clustering, we executed three independent batches of 10,000 random Louvain clustering runs. We used the Adjusted Rand Index (ARI) to measure the difference between groups from these random runs with the original groups from igraph version 1.2.11. The median ARIs were 0.92, 1.00, and 0.78 for the HPV+ exploratory set, HPV+ validation set, and HPV- exploratory set, respectively, (HPV- validation set groups were not used in this analysis), indicating small variations in the patient cluster membership. The 11- and 6-patient immune groups in the HPV+ exploratory and validation set were largely unchanged (identical in >9,200 and >8,800 out of 10,000 runs, respectively). For the HPV- exploratory set, the top five grouping results in each batch have either 30- or 32-patient groups and both groups are enriched with immune response GO terms. In summary, despite the randomness introduced in igraph version 1.3, our findings are still maintained.

### Supplementary Tables

**Supplementary Table 1.**
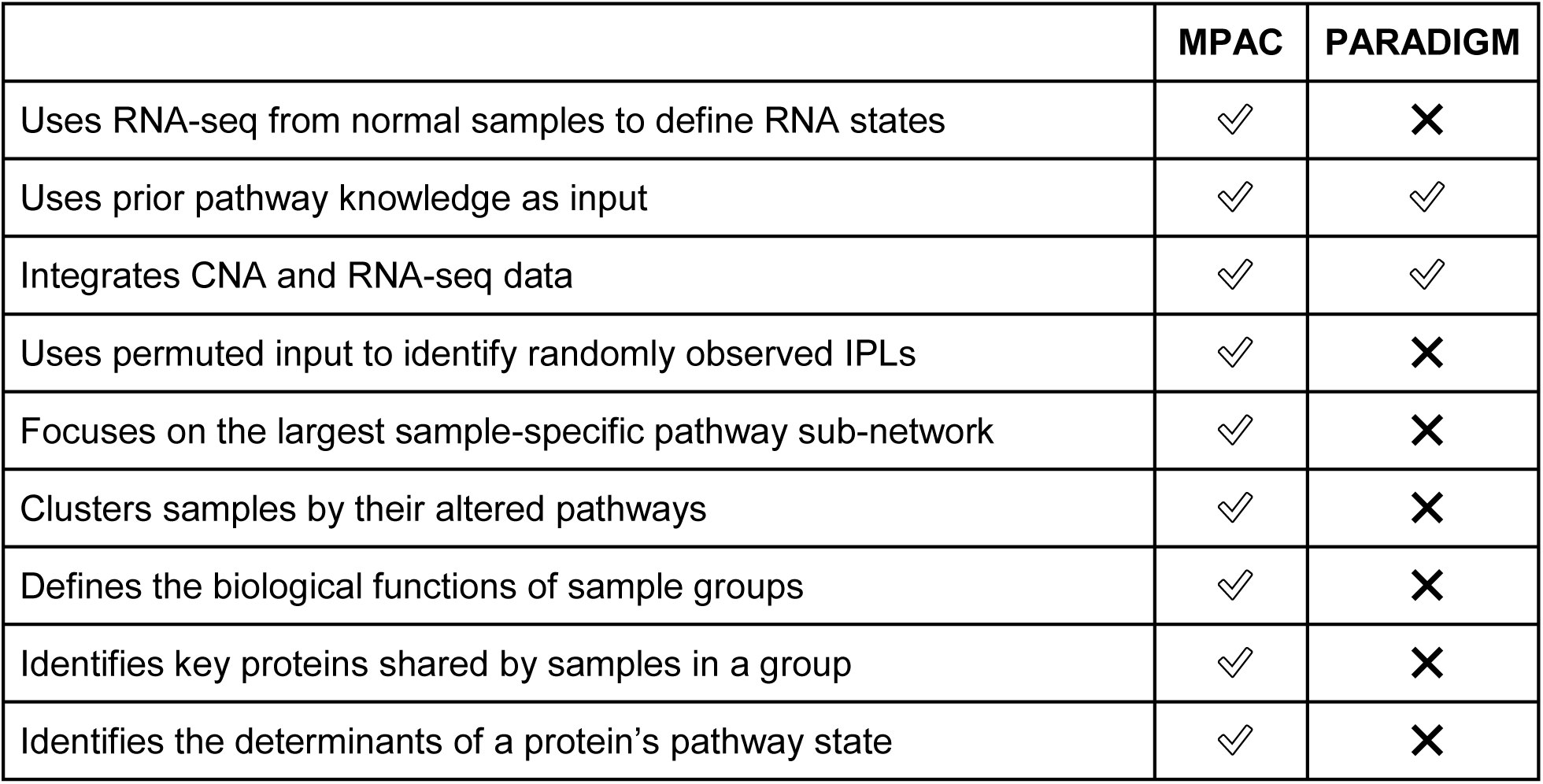
Comparison of MPAC and PARADIGM features. A check mark represents that feature is implemented and a cross mark indicates the lack of the feature.

|  | MPAC | PARADIGM |
| --- | --- | --- |
| Uses RNA-seq from normal samples to define RNA states | ✓ | ✗ |
| Uses prior pathway knowledge as input | ✓ | ✓ |
| Integrates CNA and RNA-seq data | ✓ | ✓ |
| Uses permuted input to identify randomly observed IPLs | ✓ | ✗ |
| Focuses on the largest sample-specific pathway sub-network | ✓ | ✗ |
| Clusters samples by their altered pathways | ✓ | ✗ |
| Defines the biological functions of sample groups | ✓ | ✗ |
| Identifies key proteins shared by samples in a group | ✓ | ✗ |
| Identifies the determinants of a protein's pathway state | ✓ | ✗ |

**Supplementary Table 2.**
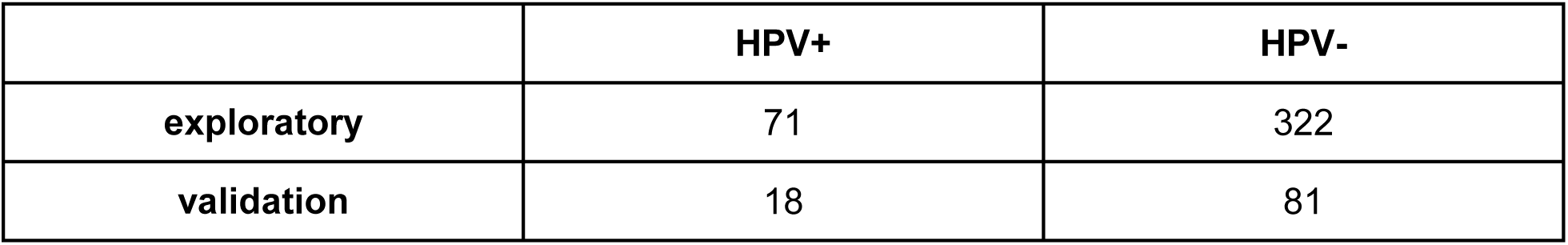
Number of patient samples in each dataset. HNSCC patient samples were stratified first by HPV subtypes and then by a random 80% and 20% split into exploratory and validation set to tune and test MPAC, respectively.

**Supplementary Table 3.**
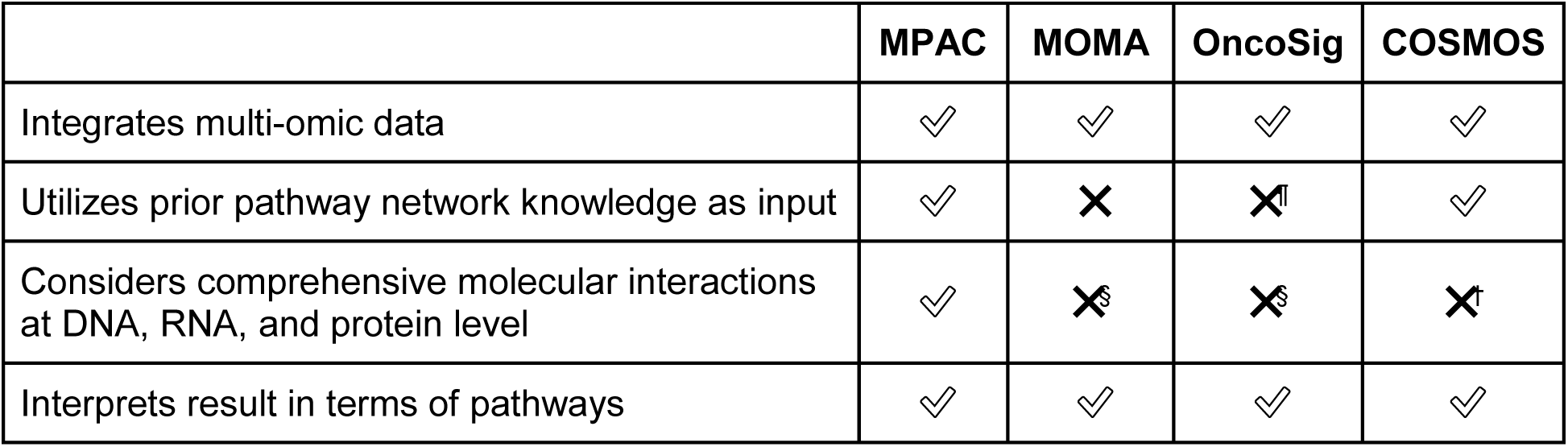
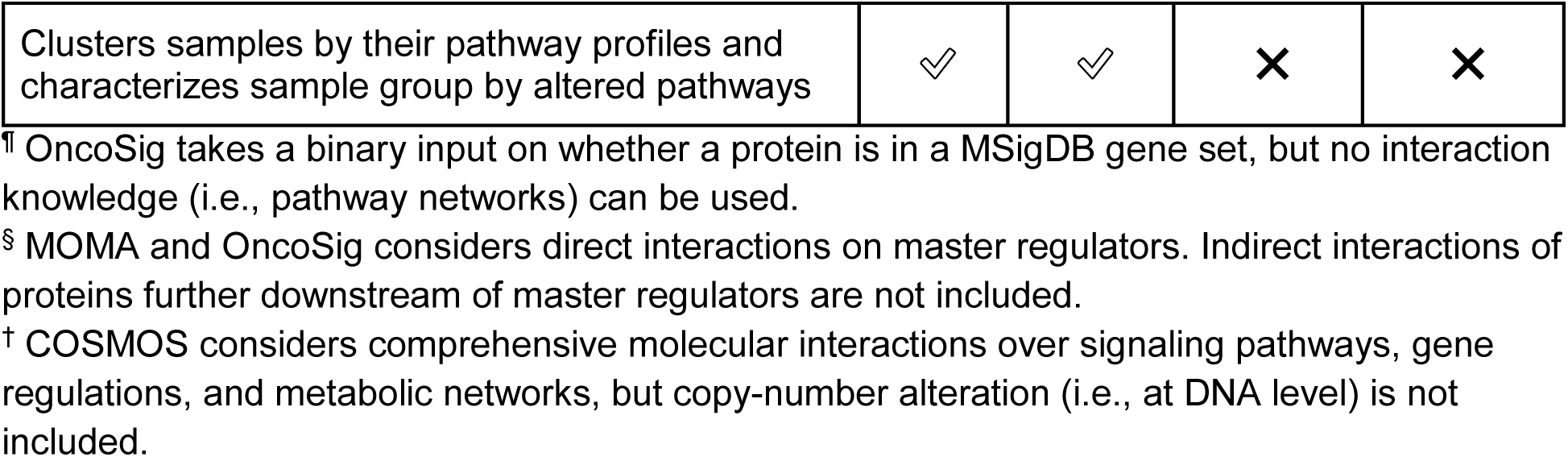
Feature comparison of MPAC with other related software. A check mark represents that feature is implemented and a cross mark indicates the lack of the feature.

|  | MPAC | MOMA | OncoSig | COSMOS |
| --- | --- | --- | --- | --- |
| Integrates multi-omic data | ✓ | ✓ | ✓ | ✓ |
| Utilizes prior pathway network knowledge as input | ✓ | ✗ | ✗ | ✓ |
| Considers comprehensive molecular interactions at DNA, RNA, and protein level | ✓ | ✗ | ✗ | ✗ |
| Interprets result in terms of pathways | ✓ | ✓ | ✓ | ✓ |
| Clusters samples by their pathway profiles and characterizes sample group by altered pathways | ✓ | ✓ | ✗ | ✗ |

<sup>¶</sup> OncoSig takes a binary input on whether a protein is in a MSigDB gene set, but no interaction knowledge (i.e., pathway networks) can be used.
<sup>§</sup> MOMA and OncoSig considers direct interactions on master regulators. Indirect interactions of proteins further downstream of master regulators are not included.
<sup>†</sup> COSMOS considers comprehensive molecular interactions over signaling pathways, gene regulations, and metabolic networks, but copy-number alteration (i.e., at DNA level) is not included.

